# Spatially variable growth responses to warming in Atlantic Sea Scallops (*Placopecten magellanicus*)

**DOI:** 10.1101/2025.10.15.682622

**Authors:** Zhengchen Zang, Rubao Ji, Deborah R. Hart, Romain Lavaud, Changsheng Chen, Kirstin Meyer-Kaiser, Siqi Li, Antonie S. Chute, Roger L. Mann

**Affiliations:** Louisiana Universities Marine Consortium, Chauvin, LA, USA; Biology Department, Woods Hole Oceanographic Institution, Woods Hole, MA, USA; National Oceanic and Atmospheric Administration Northeast Fisheries Science Center, Woods Hole, MA, USA; School of Renewable Natural Resources, Louisiana State University Agricultural Center, Baton Rouge, LA, USA; School for Marine Science and Technology, University of Massachusetts Dartmouth, New Bedford, MA, USA; Virginia Institute of Marine Science, William & Marry, Gloucester Point, VA, USA

**Author notes:** Corresponding author: Zhengchen Zang).

**Keywords:** Keyword: Atlantic sea scallop, growth rate, temperature, bioenergetics

## Abstract

Atlantic sea scallop is an important commercial species in the U.S., and its growth rate is strongly modulated by temperature. In this study, we quantified the interannual variations in scallop growth rate and thermal stress intensity in the Mid-Atlantic Bight (MAB) from 2000 to 2018. The results showed that the scallop growth rate variations were overall synchronized between the shallow (< 60 m) and deep (≥ 60 m) MAB prior to 2015. The response of growth rate to warming in 2015 and 2016 showed marked spatial heterogeneity: scallops in the shallow subregions grew more slowly in the warm years, whereas higher temperature contributed to elevated growth rates in the deep subregions. We developed a dynamic energy budget model to explain the distinct growth rate response to warming at different depths. The model results showed that, in 2016, bottom temperatures in the shallow subregions exceeded the optimal range, limiting the energy available for growth. In contrast, warming created more favorable thermal conditions in the deep habitats. This work reveals the spatiotemporal patterns of scallop growth rates in the MAB and contributes a quantitative understanding of thermal stress and the development of science-based management strategies under future warming.

## 1. Introduction

The Atlantic sea scallop (*Placopecten magellanicus*), distributed on the continental shelf between Cape Hatteras, NC, and the Gulf of St. Lawrence, is one of the most economically valuable fishery species in the U.S. (Stewart and Arnold, 1994). Although the total sea scallop landing in the U.S. fishing grounds (e.g., Mid-Atlantic, Georges Bank, and the Great South Channel) dropped dramatically from the 1970s to the mid-1990s due to overfishing (Edwards and Murawski, 1993), the implementation of a series of effective fishery management strategies has rebuilt the depleted populations since 1994 (Hart and Rago, 2006). Sea scallop stocks successfully recovered due to this species’ rapid individual growth rate and low natural mortality, and the ex-vessel revenues of sea scallop fisheries have increased to around $500 million per year in the recent decade (Hart, 2003; Coleman *et al*., 2022; NOAA, 2023).

Sea scallop growth rate is one of the most crucial biophysiological traits indicating the health condition of individuals and suitability of surrounding habitats (MacDonald and Thompson, 1985; Hart and Chute, 2009; Hodgdon *et al*., 2020; Coleman *et al*., 2021). Given the importance of scallop growth in biomass production, management plan development, and aquaculture site selection, previous studies assessed the spatiotemporal patterns of scallop growth rates in natural habitats and their associated drivers. Laboratory experiments, field measurements, and model results indicated that various factors (e.g., temperature, salinity, water depth, density, food condition, current fields, CO_2_ level, fishing activities, sediment resuspension, and parasite infection) and their joint effects modulate sea scallop growth (Grant *et al*., 1997; Pilditch and Grant, 1999a, 1999b; Hart and Chute, 2009; Inglis *et al*., 2016; Siemann *et al*., 2019; Coleman *et al*., 2022; Pousse *et al*., 2023). Among all these factors, temperature has received much attention due to its profound effects on scallop metabolic processes (e.g., clearance, ingestion, respiration, and excretion rates; MacDonald and Thompson, 1985; Shumway *et al*., 1988; Cranford and Grant, 1990; Grant and Cranford, 1991). The optimum temperature for scallop growth is between 10–15 °C, and higher temperatures limit scallop growth and survival due to net energy loss associated with decreased feeding and elevated energy costs of maintenance (Stewart and Arnold, 1994; Hart and Chute, 2004; Coleman *et al*., 2022; Zang *et al*., 2022a). Additionally, the intricate nonlinear interactions between temperature and other stressors further confound the thermal response of scallop growth. Coleman *et al*. (2021) found that in high scallop density, a negative correlation between temperature and growth could occur during some of the most productive periods when temperature was favorable to the rapid growth of individuals, indicating that high density lowered the optimal temperature for scallop growth. Food availability is another critical factor that interacts with temperature effects on scallop growth: sufficient high quality food supply may buffer the negative impacts of thermal stress on growth, while food deficiency can amplify its negative effects, especially for large individuals with higher commercial values (MacDonald and Thompson, 1985, 1986; Pilditch and Grant, 1999a; Zang *et al*., 2022a). Furthermore, laboratory experiments under various temperature and pCO_2_ conditions showed that the interactive effects of warming and ocean acidification (OA) contributed to declines in scallop growth and survivorship (Cameron *et al*., 2022).

The Mid-Atlantic Bight (MAB; Fig. 1) represents the southern extent of sea scallop habitat on the Northeast U.S. Shelf with relatively strong thermal stress (highest bottom temperature > 18 °C; du Pontavice *et al*., 2023), making its scallop populations more vulnerable to warming than those at higher latitudes. Bottom temperature in the MAB has increased at an average rate of ∼0.35°C per decade with strong spatial heterogeneity (Pershing *et al*., 2015; Saba *et al*., 2016; Kleisner *et al*., 2017; du Pontavice *et al*., 2023). Marine heatwaves in 2012 and 2016 acutely increased water temperature in the MAB and surrounding areas, significantly affecting thermal conditions and fishery yields (Mills *et al*., 2013; Pershing *et al*., 2018). Climate model projection results suggested that warming will continue in the MAB at the rate of ∼0.05 °C yr^−1^ with more heatwave events, although the strengthening of the Atlantic Meridional Overturning Circulation and a southward shift of the Gulf Stream will temporally pause warming in the next decade (Kleisner *et al*., 2017; Koul *et al*., 2024). Due to strong thermal stress in coastal waters, scallops in the MAB are mainly distributed at depths greater than 35 m (Fig. 1; Hart and Rago, 2006; Zang *et al*., 2022). Field surveys suggested that thermal stress has become the primary factor limiting sea scallop growth and survival near the southern edge of the MAB scallop habitats. For example, juvenile scallops in Virginia Beach and Delmarva did not grow into fishable sizes after fisheries closure (Virginia Beach: 1999–2001; Delmarva: 2013–2014), implying that excessive thermal stress lowered or stopped scallop growth and biomass increase (Wallace *et al*., 2018; Lee *et al*., 2019; Hart *et al*., 2020). The scope for growth model results showed that warming-induced energy deficiency limited scallop growth and deteriorated size structure with more small individuals in the southern MAB shallow areas (Zang *et al*., 2022a, 2023). The combination of high bottom temperature, rapid warming rates, and high scallop abundance makes the MAB an ideal natural laboratory for investigating the impacts of thermal stress on scallop growth.

**Fig. 1.**
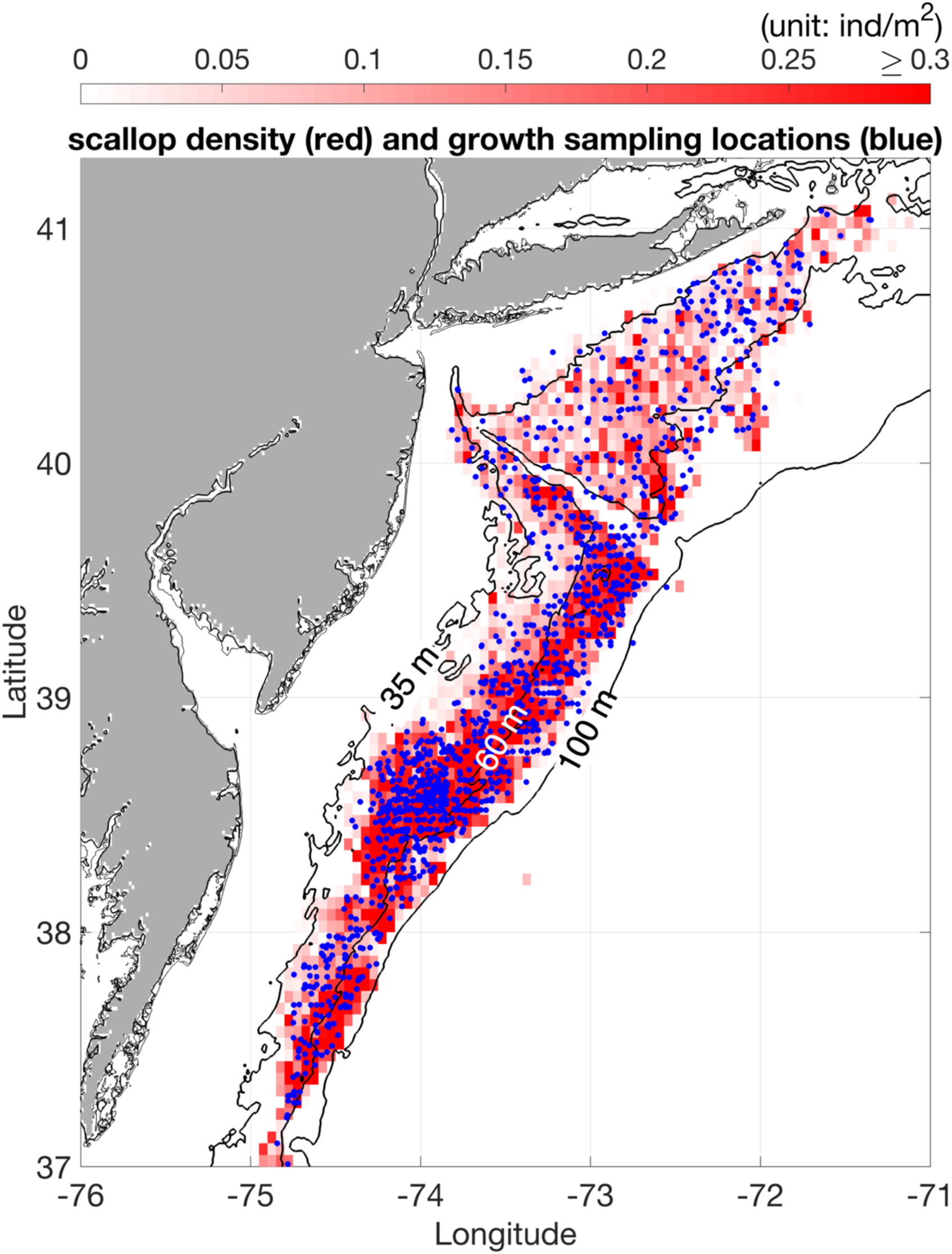
The climatology of Atlantic sea scallop density in the MAB from 2000 to 2018 (data source: NOAA sea scallop dredge survey data). The three black contour lines are 35, 60, and 100 m isobaths, respectively. The blue dots indicate locations where sea scallop shells were collected for growth measurements.

Although previous studies provided valuable information regarding the relationship between scallop growth and temperature (e.g., Parsons *et al*., 2002; Coleman *et al*., 2021), the direct evidence showing the impacts of thermal stress on sea scallop growth rate across various size groups in natural habitats is still lacking. Many studies have investigated scallop growth and its spatiotemporal variation using the von Bertalanffy growth model, which is a powerful tool in revealing scallop growth characteristics and has been widely applied to assess scallop growth variability across environmental gradients, populations, and management scenarios (Serchuk *et al*., 1979; Thouzeau *et al*., 1991; Harris and Stokesbury, 2006; Hart and Chute, 2009). However, previous von Bertalanffy growth parameter (*K* and *L*_∞_) estimations used scallop growth increment data over years. Thus, von Bertalanffy growth model results overall represent multi-year averaged scallop growth condition, and that leads to the difficulties in disentangling the impacts of warming events on shorter time scales (i.e., several months to one year). Another important knowledge gap is the spatial heterogeneity of warming effects on scallop growth. Most previous findings have highlighted the negative effects of warming on sea scallop growth and population dynamics in the MAB (Kleisner *et al*., 2017; Tanaka *et al*., 2020; Zang *et al*., 2023), but warming might also benefit the growth of those individuals in regions with low temperature below the optimum thermal conditions. The coupling/decoupling between bottom thermal conditions and scallop size structure interannual variations in the shallow/deep Delmarva rotationally closed area, respectively, also implied the spatial heterogeneity of warming effects on scallop growth (Zang *et al*., 2023).

Given the importance of growth rate in sea scallop population dynamics and its high sensitivity to temperature variations, we used long-term scallop growth data to investigate the spatiotemporal pattern of sea scallop growth rate in the MAB and its response to strong thermal stress in those warm years (e.g., 2016). The magnitude of thermal stress and its influence on scallop growth were assessed using both hydrodynamic and Dynamic Energy Budget (DEB) models. The objectives of this work were to 1) reveal the spatial patterns of sea scallop growth rate across various size groups in the MAB, 2) reconstruct the interannual variations in thermal stress intensity and scallop growth rate and examine their causal relationship, and 3) explain the spatial heterogeneity of warming effects on scallop growth from the perspective of energy balance.

## 2. Data and Methods

### 2.1 Scallop growth rate data

Sea scallop growth rate data from 2000 to 2018 were extracted from scallop shells collected during the NOAA Northeast Fisheries Science Center (NEFSC) field surveys (Fig. 1). Samples for growth rate analysis were collected at slightly less than half of the stations during each cruise, and about six samples were randomly selected at each station (Fig. 1; Hart and Chute, 2009). The scallops were scrubbed with a wire brush and shucked for laboratory analysis. Rings on the top valve of each shell were marked with a pencil, and the distance from the umbo to each of the rings was measured using calipers. The distance between two adjacent rings represents shell height growth in a year (Fig. S1). Given the difficulties in identifying the first several rings formed at early ages, only rings beyond 40 mm shell height were considered. Since the NEFSC sea scallop surveys were conducted prior to the formation of rings in fall, the distance between the two outermost rings represented shell height increment a year before, and the earlier shell height increments and their corresponding years could also be determined accordingly (Fig. S1). The partial shell increment from the last ring to the shell edge was not used in this work because it did not record the growth in a full year. To identify spatial patterns of sea scallop growth, the entire MAB was partitioned into four subregions: shallow SMAB (depth < 60 m and latitude < 39 °N), deep SMAB (depth ≥ 60 m and latitude < 39 °N), shallow NMAB (depth < 60 m and latitude ≥ 39 °N), and deep NMAB (depth ≥ 60 m and latitude ≥ 39 °N). The scallop growth rate data were categorized into eighteen size bins with 5 mm intervals from 40 mm to 130 mm based on the distance from the umbo to a ring (i.e., initial shell height at the beginning of the corresponding year; Fig. S1). Due to insufficient data for directly assessing interannual variation in scallop growth rate within each size bin, we applied z-score normalization to the measured scallop growth rates within their respective size bins and combined normalized growth rate data to reconstruct normalized growth rate interannual variation in each subregion.

### 2.2 Bottom temperature data and thermal stress intensity estimation

We extracted bottom temperature data from the outputs of the Finite Volume Community Ocean Model–Gulf of Maine Versions 3 and 4 (FVCOM–GOM3/4). FVCOM–GOM3 is a hydrodynamic model nested within the FVCOM–Global Model (Chen *et al*., 2003, 2011, 2021a). The model domain covers the entire Northeast U.S. Shelf from the Scotian Shelf to the MAB and the adjacent open ocean (Fig. S2a). The horizontal grid resolution varies from 0.5 to 10 km depending on the complexity of coastal geometry and steep bottom topography. The model grid is discretized into 45 sigma layers in the vertical direction. Measured temperature, salinity, and current profiles are assimilated into the model results to enhance the quality of its outputs (Chen *et al*., 2009, 2021b). The modeled water temperature products have been well-calibrated and widely used in previous studies (Chen *et al*., 2011, 2021b; Li *et al*., 2015; Zang *et al*., 2022b). Since the simulations of FVCOM-GOM3 were terminated in 2016, bottom temperature data after 2016 were extracted from FVCOM-GOM4. FVCOM-GOM4 is an updated version of the FVCOM-GOM3 with the same spatial coverage (Fig. S2b), and its grid resolution is higher in nearshore areas (e.g., Penobscot Bay, Boston Harbor, and Narragansett Bay) to better simulate coastal processes. The hourly hydrodynamic model outputs were downloaded from the data server of the University of Massachusetts Dartmouth (http://fvcom.smast.umassd.edu) and converted to daily average results.

To assess thermal stress intensity in the MAB, the modeled bottom temperature data were spatially interpolated to the locations of scallop samples using the corresponding shell height increment year determined in Section 2.1. Since the optimum temperature for sea scallop growth is below 15 °C (Coleman *et al*., 2022; Zang *et al*., 2022a), we estimated Degree Days > 15 °C (DD hereafter) to represent thermal stress intensity:

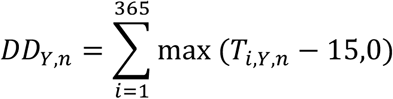

where *T*_*i*,*Y*,*n*_ is the daily averaged bottom temperature on Julian day *i* of year *Y* for sample *n*. Since sea scallops in the MAB lay their rings when temperatures peak in the fall season (October to December; Chute *et al*., 2012), DD for year *Y* was estimated using temperature data from Nov 1^st^ of year *Y*-1 to Oct 31^st^ of year *Y* to ensure that the temporal coverage of DD matches that of shell height increment (Section 2.1).

### 2.3 Sea scallop DEB model

We developed a sea scallop bioenergetic model based on the Dynamic Energy Budget theory (Kooijman, 2010). The DEB model simulates energy fluxes through four state variables: 1) the structural volume *V*; 2) the reserve *E*, 3) the energy allocated to maturity and reproduction *E*_*R*_, and 4) the energy used to create gametes *E*_*Go*_ (Fig. 2). Temperature and food availability are the forcing variables. Energy from food is assimilated into the reserve compartment, and a fixed fraction (*κ*) of energy flux from the reserve is allocated to support somatic maintenance and structure growth. The rest of the energy (1-*κ*) is used for maturity maintenance, maturation, and reproduction. The primary DEB model parameters for *P. magellanicus* were retrieved from the AmP collection (Lavaud and Kooijman, 2020). We adopted the DEB model structure in Lavaud *et al*. (2014, 2017, 2019, 2021) to simulate sea scallop growth and its response to temperature variation, and this DEB model structure has been successfully applied to simulate the growth and reproduction of other bivalve species (e.g., king scallop and eastern oyster). Readers are referred to Lavaud *et al*., (2014, 2017) for more detailed information regarding the DEB model structure. Due to the distinct optimum temperatures for sea scallop feeding and maintenance (Shumway *et al*., 1988), we applied different parameters in the Arrhenius relationships (i.e., *T*_*AL*_ and *T*_*AH*_) for energy assimilation and consumption to better resolve the effects of warming on scallop growth. All model equations and parameters are listed in Tables. S1 and S2. Daily bottom temperature climatology in the entire MAB scallop habitats was estimated and used to drive the DEB model. The simulation was initiated with a scallop shell height of 40 mm (year 0) and lasted 14 years. The modeled shell height was compared with observed shell height in the MAB based on von Bertalanffy growth parameters estimated by Hart and Chute (2009) (*K* = 0.508 and *L*_∞_ = 133.3 *mm*). The simulated tissue weight and gonad weight were compared with the data using the log_10_-log_10_ relationships between shell height and these variables in Langton *et al*. (1987).

**Fig. 2.**
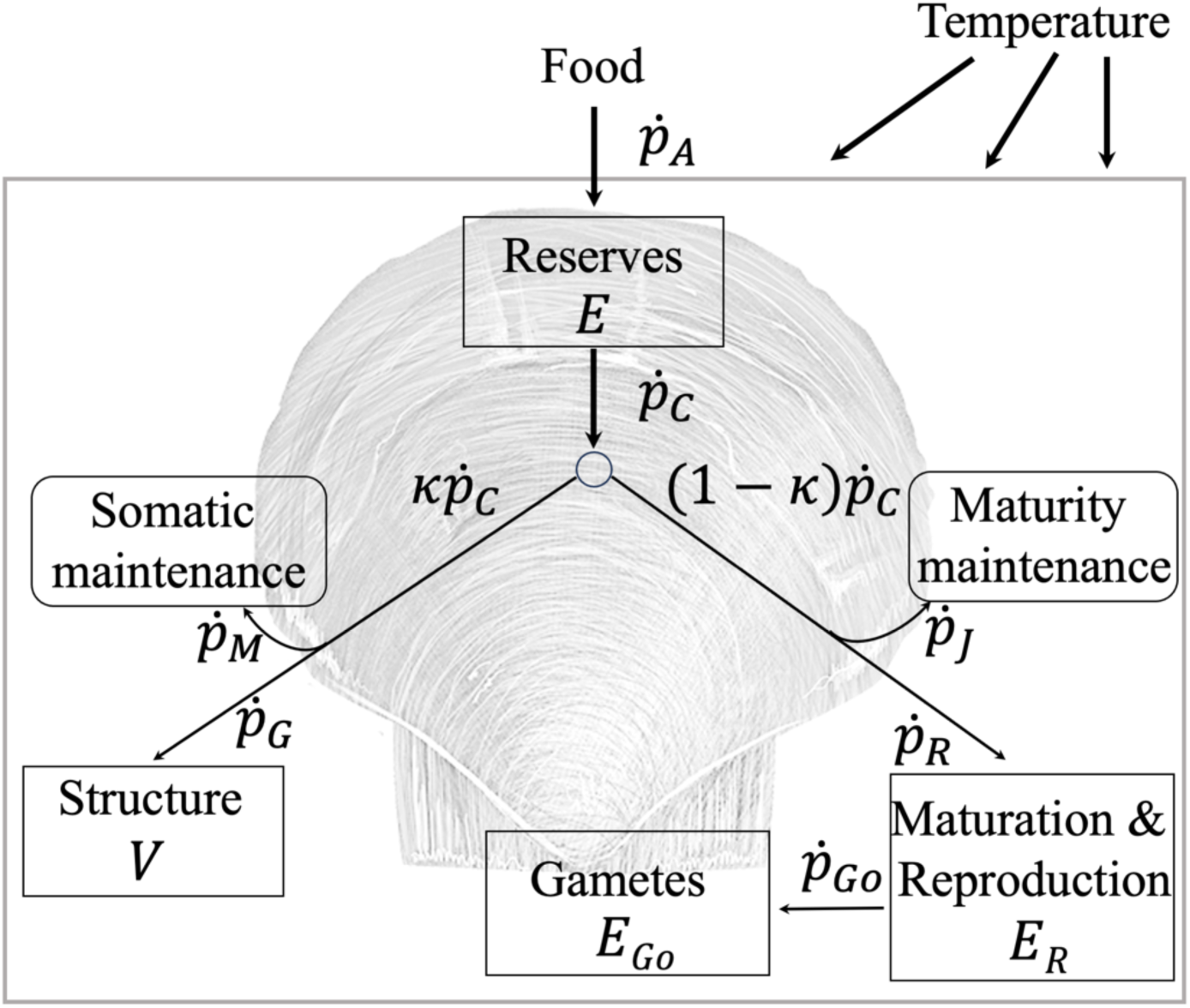
Conceptual scheme of the DEB model applied to sea scallop. Four state variables are Reserves (*E*), Structure (*V*) and Maturity & reproduction (*E*_*R*_), and Gametes (*E*_*Go*_) in rectangular boxes. *ṗ*_*A*_, *ṗ*_*C*_, *ṗ*_*M*_, *ṗ*_*G*_, *ṗ*_*J*_, *ṗ*_*R*_, and *ṗ*_*Go*_ are energy fluxes estimated in the model (see Table S1 in supplementary material for more detailed information). Arrows indicate energy flux directions.

To quantitatively estimate warming effects on sea scallop growth and its spatial heterogeneity, we designed sensitivity tests driven by daily bottom temperature climatology and bottom temperature in the warm year 2016 in the four subregions. All other inputs and parameters were the same to ensure that the differences between model results only resulted from temperature inputs. The simulation period of sensitivity tests was one year, and we used four initial sea scallop shell heights (i.e., 40 mm, 60 mm, 80 mm, and 100 mm) to assess the impacts of warming on scallop growth in different size groups.

## 3. Results

### 3.1 Spatial pattern of scallop growth rate in the MAB

The median scallop growth rate decreased gradually from ∼ 37 mm yr^−1^ in the smallest size bin (40–45 mm) to 6 mm yr^−1^ in the largest size bin (125–130 mm) (Fig. 3). In both the SMAB and the NMAB, scallop growth rates across almost all size bins were significantly higher in the shallow subregions (Fig. 3). Growth rate differences between the shallow and deep subregions were most pronounced in the 60–80 mm size bins, reaching ∼ 5–9 mm yr^−1^; whereas differences for smaller and larger scallops were generally below 4 mm yr^−1^. Unlike the significant growth rate difference between shallow and deep areas, the growth rate differences across latitudes within the same depth range were much smaller (Fig. S3). In the shallow subregions, individuals between 70 and 100 mm grew 1–3 mm yr^−1^ faster in the NMAB, while only marginal differences were detected in other size groups (Fig. S3a). Growth rates in the deep SMAB and the deep NMAB were quite comparable across all size bins (Fig. S3b). Overall, sea scallop growth rates showed a much stronger and consistent gradient across depths than across latitudes in the MAB, with higher growth rates observed in shallow areas in all size groups.

**Fig. 3.**
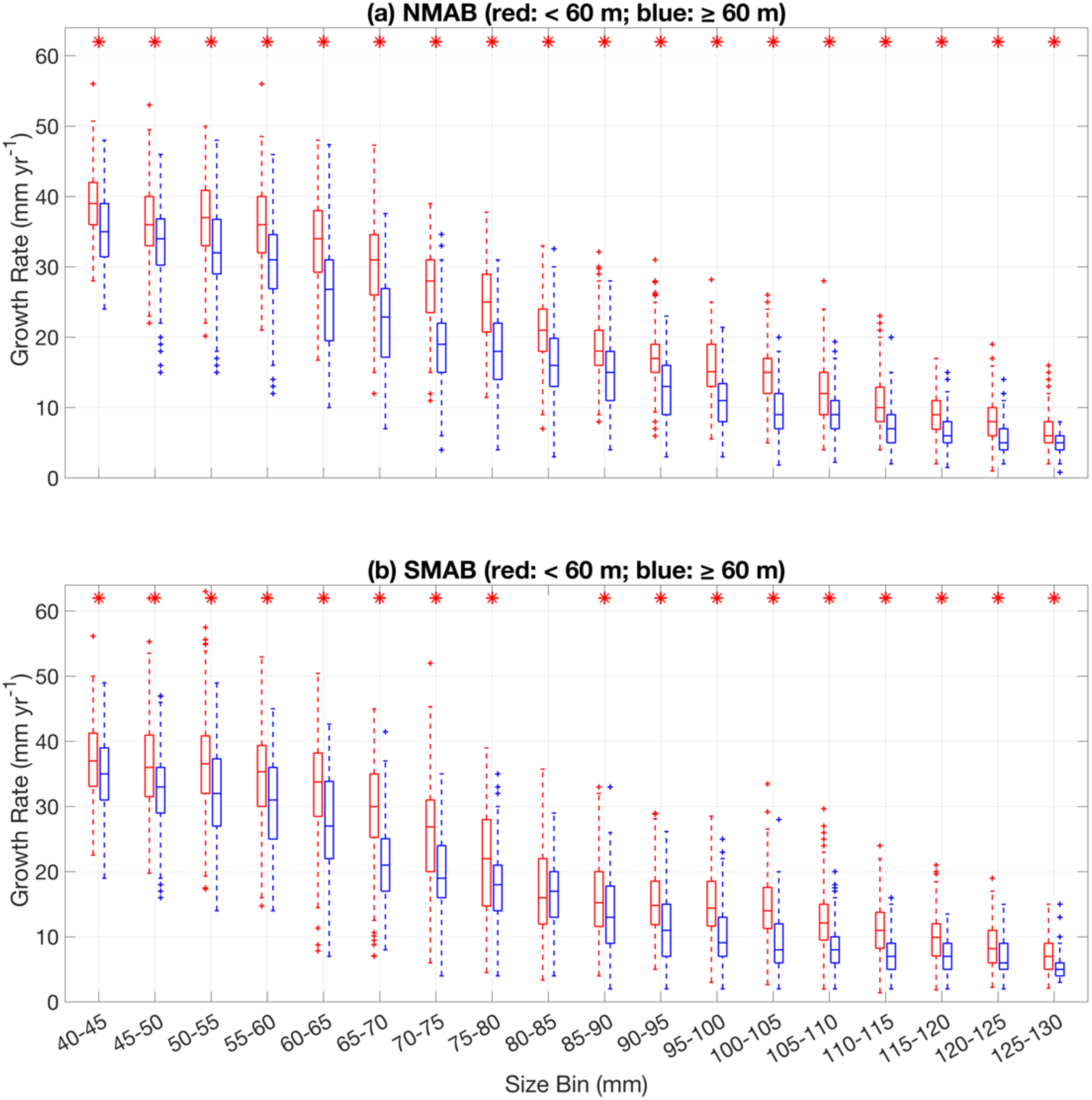
Scallop growth rates across 18 size bins (x-axis) in the northern MAB (NMAB; panel a) and the southern MAB (SMAB; panel b). Red and blue colors indicate data in the shallow and the deep subregions, respectively. Mann–Whitney U tests were performed for each size bin to assess statistical significance. Red asterisks indicate significantly higher values in the shallow subregions.

### 3.2 Interannual variations in normalized scallop growth rate and thermal stress

The significant positive correlations between normalized growth rates in the two shallow subregions (shallow SMAB – shallow NMAB: r = 0.76, p < 0.05) and the two deep subregions (deep SMAB – deep NMAB: r = 0.64, p < 0.05) indicated that the interannual variations in scallop growth rates were synchronized within the same depth range (Fig. 4). In contrast, no significant correlation was detected between the shallow and deep subregions within the NMAB (shallow NMAB – deep NMAB: *r* = 0.41, *p* > 0.05) or the SMAB (shallow SMAB – deep SMAB: *r* = 0.24, *p* > 0.05) from 2000 to 2018. Interestingly, the decoupling of normalized growth rates across depths did not last throughout the entire study period. Correlation analyses using 14-year moving time windows indicated that the interannual variations in normalized growth rates were positively correlated between the shallow and deep subregions prior to 2015 (Table 1), while the correlation coefficients gradually declined and became statistically insignificant after 2015 (Table 1). The time series showed that normalized growth rates in the shallow subregions gradually declined below 0 after 2015 (Fig. 4a and 4c), whereas growth rates in the deep subregions remained high (deep NMAB: 0.3–0.6; deep SMAB: 0.7–1.0; Fig. 4b and 4d). The contrasting post-2015 growth patterns between shallow and deep subregions contributed to the insignificant correlations when the time window included years after 2015. We also estimated the interannual variations in normalized growth rates based on only small (≤ 80 mm) and only large (> 80 mm) individuals (Fig. S4). The results showed similar trends to those across all size classes, indicating overall consistent spatiotemporal patterns in scallop growth rates in different size groups. In addition, we estimated scallop growth rate anomalies in the four subregions using observational data and simulated growth rate based on von Bertalanffy growth curve in Hart and Chute (2009) to exclude depth and latitude effects (Supporting Text in Supplementary Material). The results were similar to our normalized growth rate interannual variations with elevated/decreased growth rates in the deep/shallow subregions after 2015, respectively (Fig. S5), confirming the spatial heterogeneity of post-2015 scallop growth patterns.

**Fig. 4.**
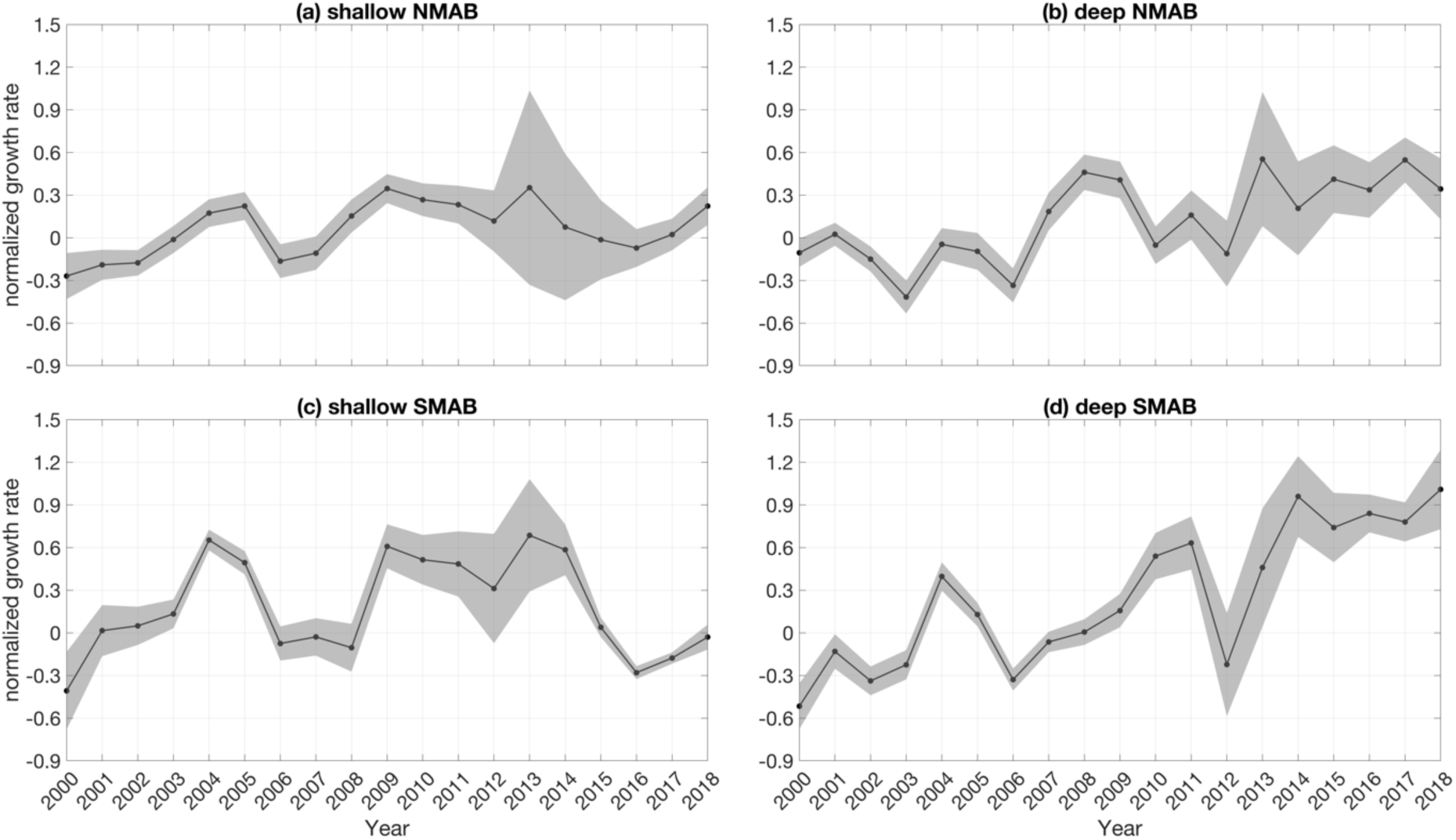
Normalized scallop growth rate interannual variations in four subregions from 2000 to 2018 (panel a: shallow NMAB; panel b: deep NMAB; panel c: shallow SMAB; panel d: deep SMAB). The grey envelops represent 95% confidence intervals.

**Table 1.** Normalized scallop growth rate interannual correlations within 14-yr time windows between the shallow and deep MAB subregions (*: *p* < 0.05; **: *p* < 0.01).

|  | 14-yr Time Window |  |  |  |  |  |
| --- | --- | --- | --- | --- | --- | --- |
|  | 2000-2013 | 2001-2014 | 2002-2015 | 2003-2016 | 2004-2017 | 2005-2018 |
| **NMAB (shallow vs. deep)** | 0.55* | 0.54* | 0.49 | 0.34 | 0.20 | 0.25 |
| **SMAB (shallow vs. deep)** | 0.82** | 0.76** | 0.57* | 0.24 | 0.08 | -0.01 |

Interannual variations in DD (i.e., thermal stress intensity) in the four subregions stayed low and fluctuated between 0 and 100 °C·d before 2015 (Fig. 5). In 2015 and 2016, DD in shallow subregions increased to 150–210 °C·d, which was followed by a gradual decrease after 2017 (Fig. 5a and 5c). In the deep SMAB, DD peaked in 2015 and 2016, but its magnitude was lower than that in the shallow subregions (2015: 140 °C·d; 2016: 121 °C·d; Fig. 5d). DD in the deep NMAB was not significantly higher in 2015 and 2016 compared to other years (Fig. 5b), suggesting that the scallops in the deep NMAB did not experience strong thermal stress likely due to its higher latitude and greater depth.

**Fig. 5.**
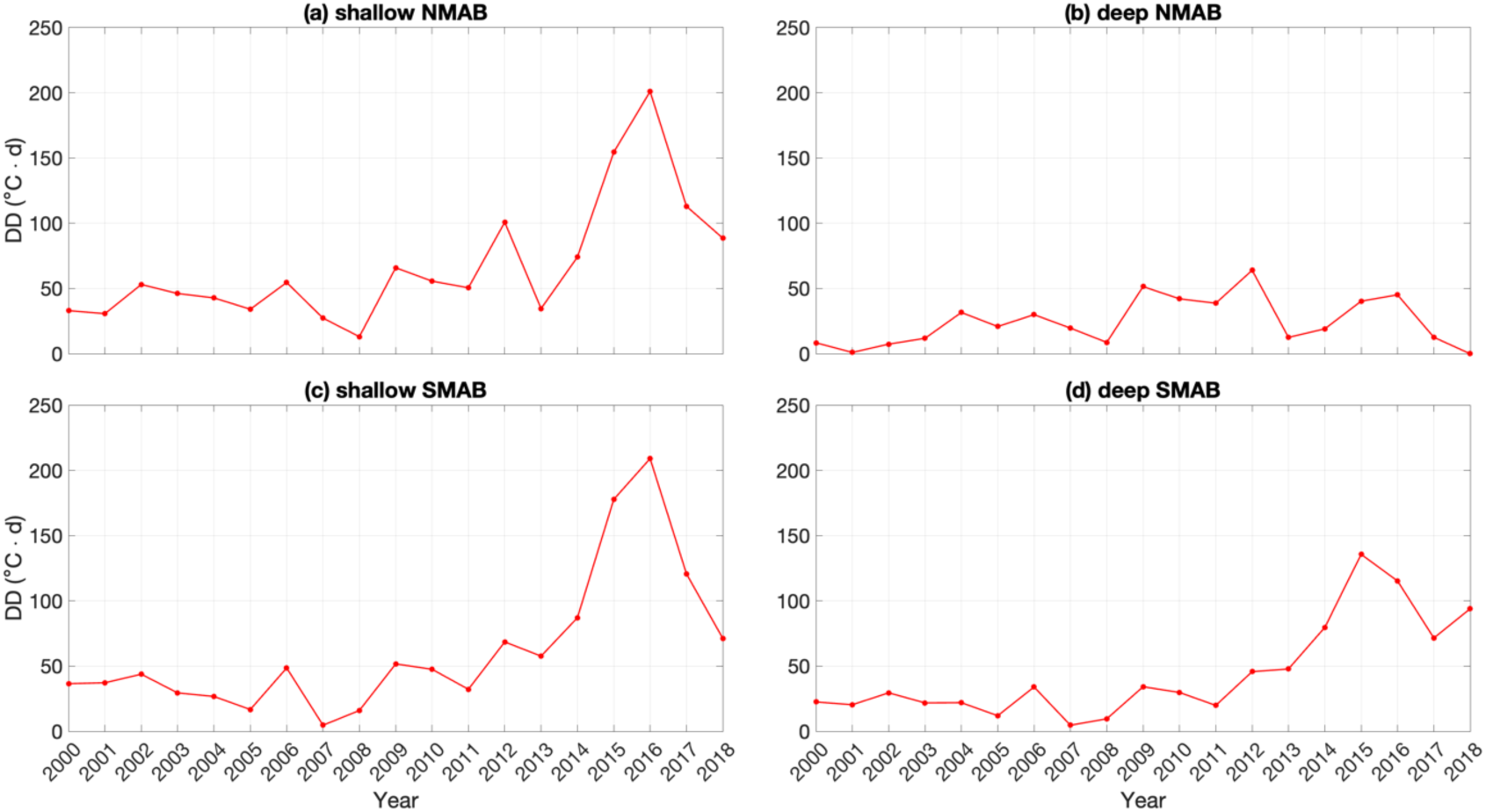
Degree days (DD, unit: °C ·d) interannual variations in the four subregions from 2000 to 2018 (panel a: shallow NMAB; panel b: deep NMAB; panel c: shallow SMAB; panel d: deep SMAB).

### 3.3 DEB model results

To evaluate the performance of our DEB model in simulating sea scallop growth in the MAB, we compared modeled sea scallop shell height, dry tissue weight, wet tissue weight, dry gonad weight, and wet gonad weight with previous statistical results based on laboratory measurements (Langton *et al*., 1987; Hart and Chute, 2009) (Fig. 6). Both the DEB model results and von Bertalanffy growth curve in Hart and Chute (2009) showed that scallop shell height in the MAB reach an asymptote at 133 mm within eight years (Fig. 6a). Modeled dry and wet tissue weights were comparable to the statistical results of Langton *et al*. (1987), increasing from approximately 1 to 20 g and 5 to 160 g, respectively (Fig. 6b and 6c). The seasonal fluctuations of dry and weight tissue weights were driven by reproduction (Fig. 6b and 6c). Dry and wet gonad weights decreased dramatically during two spawning events per year and recovered gradually after spawning (Fig. 6d and 6e). The fluctuation magnitude of modeled dry gonad weight increased gradually with age from almost 0 g in year 0 to 3 g after year 8 (minimum dry gonad weight = 0.8 g; maximum dry gonad weight = 3.8 g) (Fig. 6d). Wet gonad weight fluctuated within a wider range between 7 and 25 g after scallops reaching their maximum size in year 8 (Fig. 6e).

**Fig. 6.**
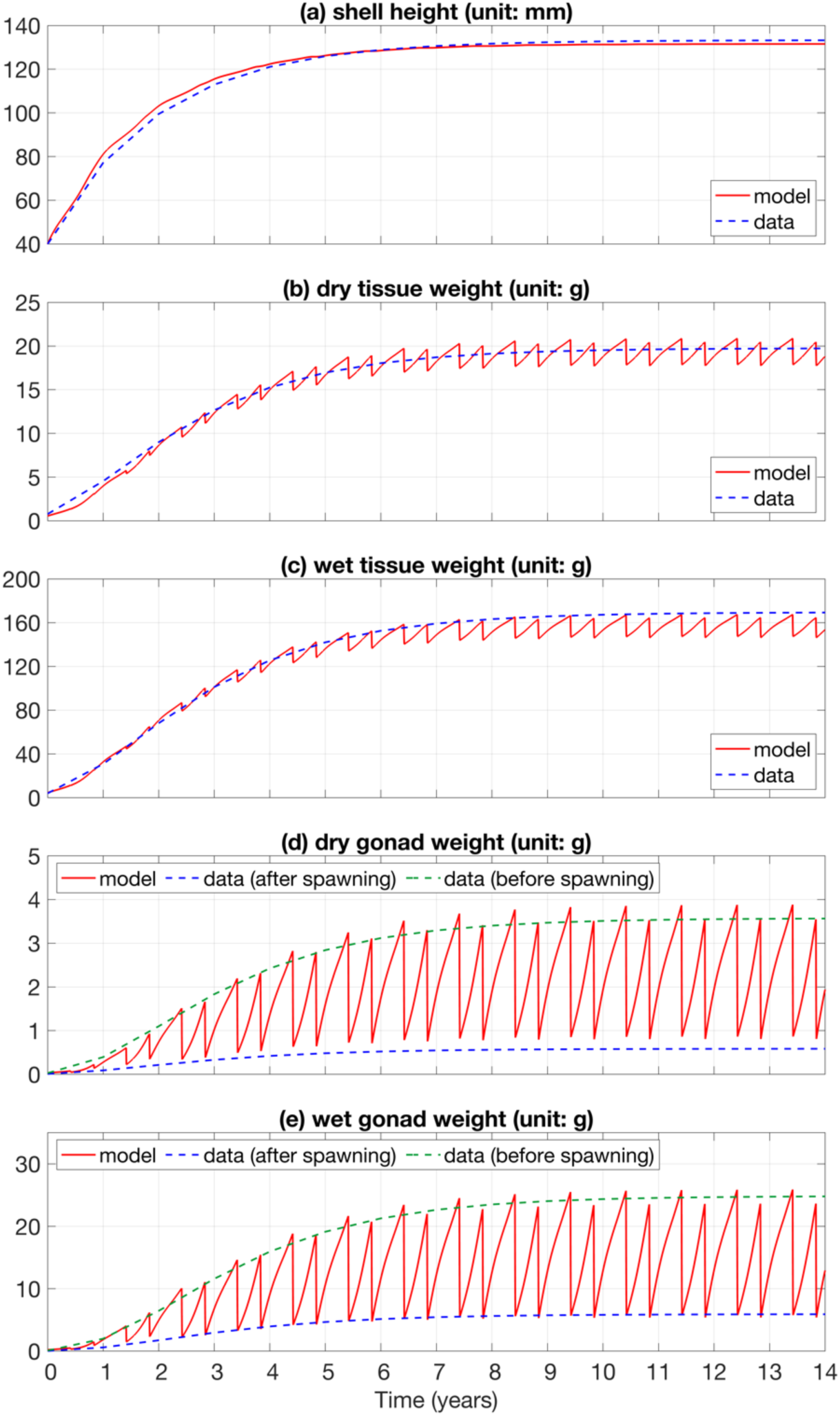
Simulated shell height (SH; a), dry tissue weight (DTW; b), wet tissue weight (WTW; c), dry gonad weight (DGW; d), and wet gonad weight (WGW; e) (red solid lines) in comparison with growth curves based on laboratory-measured data in Hart and Chute (2009) and Langton *et al*. (1987) (dashed lines). The dashed line in panel a is based on von Bertalanffy growth parameters estimated by Hart and Chute (2009) (*K* = 0.508 and *L*(= 133.3 *mm*). From panels b and e, the measured dry/wet tissue weight and gonad weight are based on log_10_-log_10_ relations in Langton *et al*. (1987) (log_10_(DTW)=−4.42+2.69·log_10_(SH); log_10_(WTW)=−4.40+3.12·log_10_(SH); log_10_(DGW_before_spawning_)=−8.01+4.03·log_10_(SH); log_10_(DGW_after_spawning_)=−7.52+3.43·log_10_(SH); log_10_(WGW_before_spawning_)=−8.40+4.61·log_10_(SH); log_10_(WGW_after_spawning_)=−8.13+4.19·log_10_(SH)).

Fig. 7 shows the modeled scallop shell height (initial shell height = 40 mm; blue lines) over one year based on bottom temperature climatology (red solid lines) and 2016 bottom temperature (red dashed lines) in the four subregions. Bottom temperature in the MAB exhibited strong seasonality with higher temperature in early fall (October–November) and lower temperature from March to May (Fig. 7). Bottom temperatures in 2016 exceeded the historical climatology by 1–2 °C during most of the year (red lines in Fig. 7). In the shallow SMAB, the bottom temperature difference between the climatology and year 2016 reached more than 5 °C in early September and October (Fig. 7c). In the shallow subregions, scallop shell height was predicted to grow faster in 2016 than under climatological temperature in the first eight months because temperature in 2016 was closer to the optimum temperature (< 15 °C) for scallop growth (Fig. 7a and 7c). From September to November, bottom temperature exceeding 15 °C in 2016 almost halted scallop growth, whereas shell height continued to increase under climatological temperature, resulting in approximately 5 mm greater shell height by the end of the simulation (Fig. 7a and 7c). In the deep subregions, although bottom temperature in 2016 was overall higher than the climatology, it barely exceeded 15 °C and facilitated faster scallop growth (Fig. 7b and 7d). The modeled scallop shell height in 2016 was approximately 4 mm higher than under the climatology conditions by the end of the one-year simulation (Fig. 7b and 7d). The sensitivity tests based on 60-, 80-, and 100-mm initial shell heights showed similar spatial patterns, although larger individuals in the deep subregions benefited less from warming (Figs. S6–S8). In general, the DEB model results reflected the same spatial heterogeneity in warming effects on scallop growth as observed in the data, with decreased growth rates in the shallow subregions and increased rates in the deep areas.

**Fig. 7.**
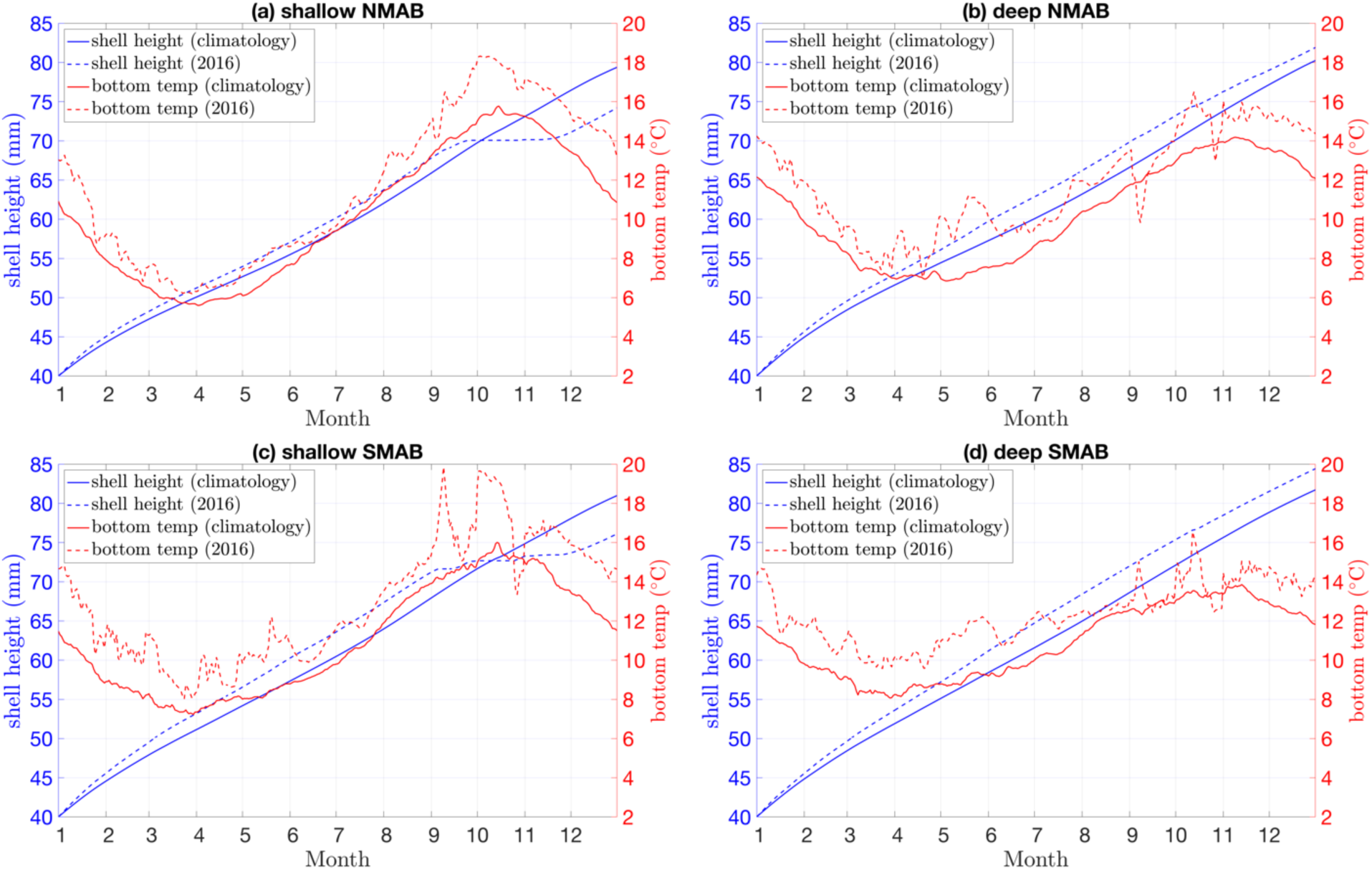
Annual shell height growth (blue) and the corresponding bottom temperature (red) in the four subregions (panel a: shallow NMAB; panel b: deep NMAB; panel c: shallow SMAB; panel d: deep SMAB). The initial scallop shell height is assigned as 40 mm. Solid and dashed lines are the results based on bottom temperature climatology and bottom temperature in 2016, respectively.

To examine the drivers of scallop growth spatial heterogeneity under warming from a bioenergetics perspective, we analyzed three energy flux in the model directly related to scallop growth: energy allocated for somatic maintenance and structure growth (*κṗ*_*C*_; Fig. 8), somatic maintenance flux (*ṗ*_*M*_; Fig. 9), and allocation flux to growth (*ṗ*_*G*_; Fig. 10). *κṗ*_*C*_ in the shallow subregions increased gradually from 50 J d^−1^ in January to ∼270 J d^−1^ in September, and its value in 2016 was 10–50 J d^−1^ higher than the climatological results (Fig. 8a and 8c). After September, *κṗ*_*C*_ became stable without a strong difference between the two temperature scenarios (Fig. 8a and 8c). In the deep subregions, *κṗ*_*C*_ increased during the entire year, and its value in 2016 was always higher than climatological results, reaching ∼400 J d^−1^ by the end of the simulation (Fig. 8b and 8d). The fluctuations of *κṗ*_*C*_ in the MAB were overall positively correlated with bottom temperature, indicating that warming enhanced energy flux for somatic maintenance and structure growth (Fig. 8). The spatiotemporal patterns of *ṗ*_*M*_ in the four subregions were very similar to those of *κṗ*_*C*_, and their comparable ranges (*κṗ*_*C*_: 50–420 J d^−1^; *ṗ*_*M*_: 50–400 J d^−1^) suggested that a substantial portion of the allocated energy was used for somatic maintenance (Fig. 9). Compared with *κṗ*_*C*_ and *ṗ*_*M*_, *ṗ*_*G*_ was considerably lower ranging from 0 to 16 J d^−1^ (black lines in Fig. 10). *ṗ*_*G*_ increased moderately from 7 J d^−1^ in January to 15 J d^−1^ in August when temperature was below 14 °C, whereas further temperature increase exceeding 15 °C led to the negative correlation between *ṗ*_*G*_ and temperature after August (Fig. 10). In the shallow subregions, *ṗ*_*G*_ in 2016 decreased to zero when bottom temperature was higher than 16 °C between September and November, whereas the climatological *ṗ*_*G*_ during the same period fluctuated around 14 J d^−1^ (Fig. 10a and 10c). In the deep subregions, although *ṗ*_*G*_ in 2016 dropped sharply below 6 J d^−1^ when temperature exceeded 16 °C in mid-October, this decline only lasted a few days before recovering to 15 J d^−1^ (Fig. 10b and 10d).

**Fig. 8.**
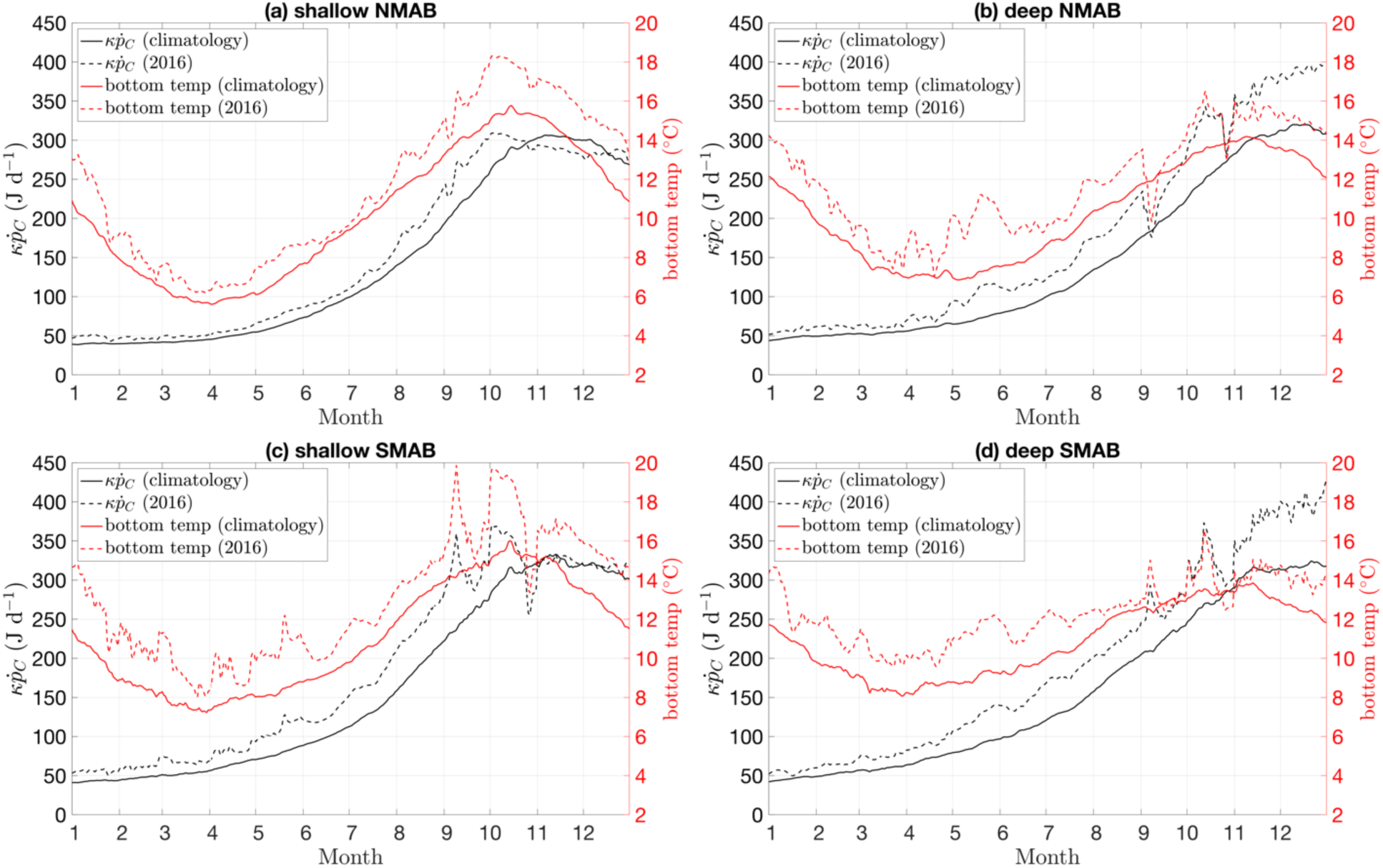
Annual energy flux for somatic maintenance and structure growth (*κṗ*_*c*_; black) and the corresponding bottom temperature (red) in the four subregions. Solid and dashed lines are the results based on bottom temperature climatology and bottom temperature in 2016, respectively.

**Fig. 9.**
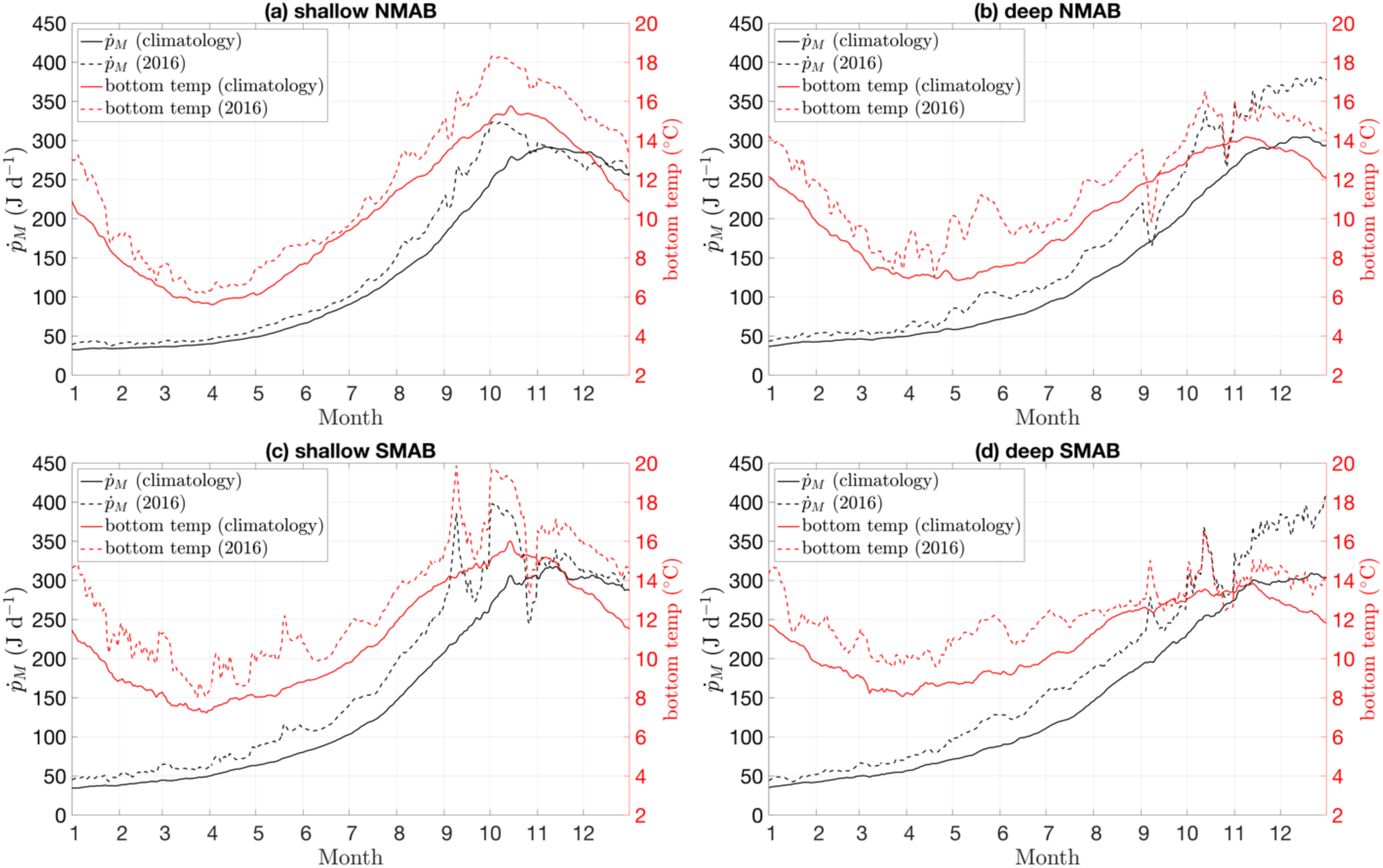
Annual somatic maintenance flux (*ṗ*_*M*_; black) and the corresponding bottom temperature (red) in the four subregions. Solid and dashed lines are the results based on bottom temperature climatology and bottom temperature in 2016, respectively.

**Fig. 10.**
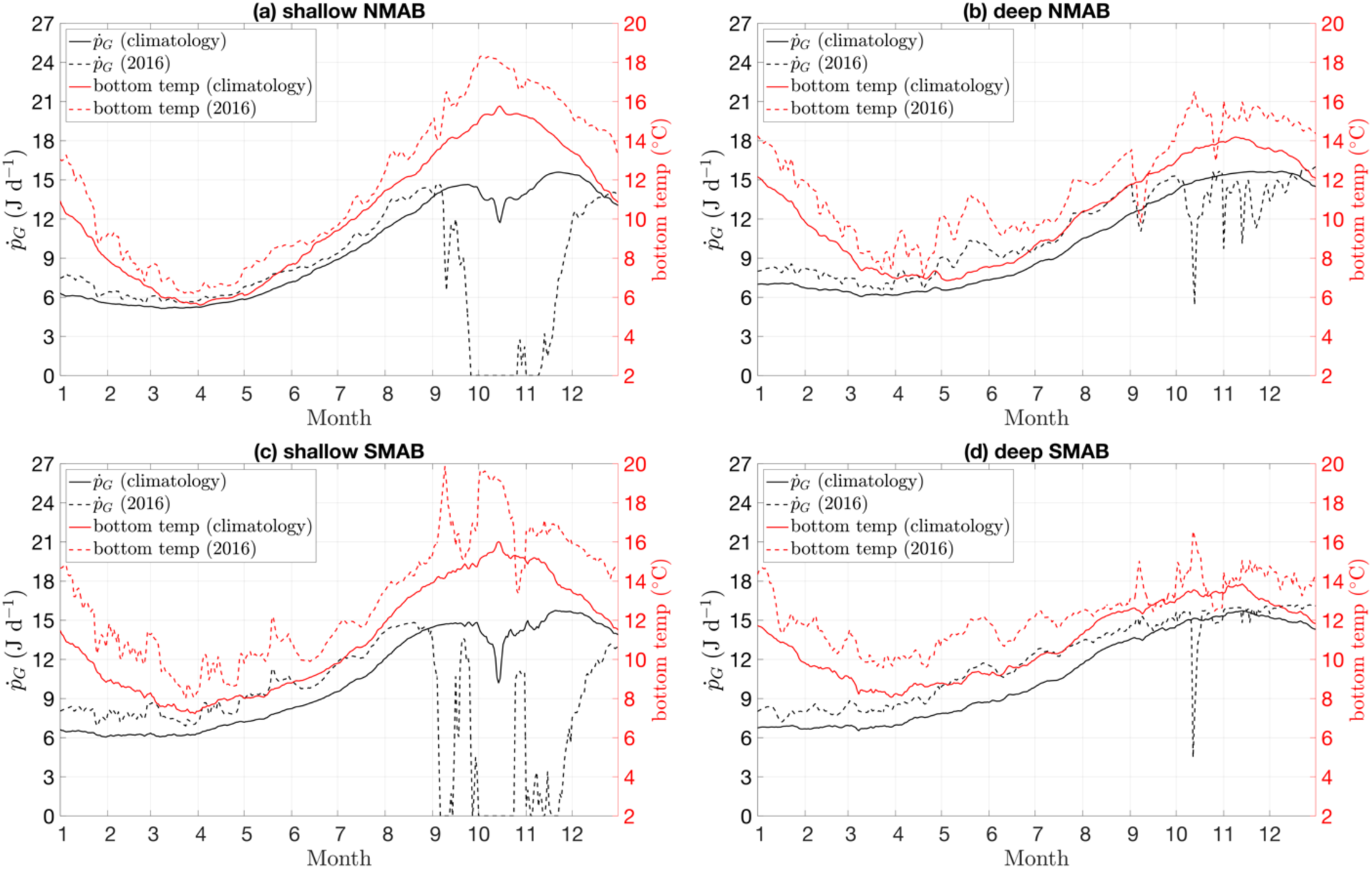
Annual allocation flux to growth (*ṗ*_*G*_; black) and the corresponding bottom temperature (red) in the four subregions. Solid and dashed lines are the results based on bottom temperature climatology and bottom temperature in 2016, respectively.

## 4. Discussion

### 4.1 Defining and quantifying thermal stress on sea scallop growth

The sea scallop is a cold-water bivalve species, and its physiological rates (e.g., feeding, growth, respiration, and mortality), biogeographic distribution, population size structure, and commercial value are very sensitive to temperature variations (MacDonald *et al*., 1987; Tanaka *et al*., 2020; Cameron *et al*., 2022; Zang *et al*., 2023). Therefore, appropriately defining and quantifying thermal stress intensity is critically important for developing a causal linkage between thermal conditions and scallop population dynamics. Mean temperature across various temporal scales has been widely applied to examine the impacts of thermal conditions on scallop growth and spatial distribution in natural habitats (Torre *et al*., 2019; Hodgdon *et al*., 2020; Tanaka *et al*., 2020). Although mean temperature can overall represent thermal environments in scallop habitats during a certain period, it smooths out high frequency variability that also plays a critical role in regulating sea scallop physiology. Laboratory experiment results indicated that temperature variation between 6 and 15 °C contributed to faster scallop soft tissue (excluding adductor) growth (Pilditch and Grant, 1999a). It should be noted that the temperature fluctuation range in Pilditch and Grant (1999a) was determined based on field measurements in Nova Scotia and did not exceed the optimum temperature for scallop growth. In the MAB where higher bottom temperatures occur, temperature fluctuations might amplify thermal stress and limit scallop growth due to prolonged exposure to unfavorable thermal conditions (Silina, 2023). Additionally, mean temperature fails to capture the phenological characteristics of temperature variability, which is also important in regulating scallop growth. Our DEB model results suggested that warming in the cold season accelerated scallop growth in the entire MAB because increased temperature was closer to the optimum temperature range. In contrast, temperature increase in the warm season lead to stronger thermal stress and was detrimental to scallop growth in the shallow subregions (Figs. 7 and 10). Thus, mean temperature obviously is not the most appropriate metric for quantifying thermal stress intensity, and that might explain the lack of significant correlation between mean temperature and scallop growth rates in some natural habitats (Hodgdon *et al*., 2020).

Other metrics have also been developed to quantify thermal stress intensity for stenothermal species (e.g., coral reefs and seagrass), and the most widely used ones are DD applied in this study and Degree Heating Weeks (DHW) (Liu *et al*., 2003, 2006; Ojea *et al*., 2008; Frieler *et al*., 2013; Strydom *et al*., 2020). DD and DHW estimate the linear accumulation of thermal stress over certain periods and explicitly include information regarding temperature, phenology of warming, and the biophysiological features of target species (temperature threshold for DD and DHW estimations). Although DD and DHW better measure thermal stress intensity than mean temperature, these metrics are most likely to be applicable to the organisms whose metabolic rates vary linearly with temperature because they don’t take the nonlinear thermal response of organisms into account (Ralston *et al*., 2014; Strydom *et al*., 2020). For example, both constant temperature of 16 °C lasting 210 days and 21 °C lasting 35 days result in equivalent DD value for sea scallops ((16 - 15) × 210 = 210 °C·d; (21 - 15) × 35 = 210 °C·d). The former scenario probably only imposes sublethal effects on scallops through lowering their growth rates (Fig. 5c), whereas the latter scenario leads to mass mortality (Stewart and Arnold, 1994; Hart and Chute, 2004). Therefore, it is reasonable to speculate that thermal stress associated with short-term acute warming is stronger than that induced by long-term moderate warming, even though their corresponding DD or DHW are comparable. To better quantify thermal stress intensity and its relationship with sea scallop growth and mortality, metrics like DD and DHW can be improved by incorporating the detailed information regarding the nonlinearities of scallop metabolic rates, thermal tolerance, and plasticity under various thermal conditions.

High quality bottom temperature data with sufficient spatiotemporal resolutions are essential for assessing thermal stress and its effects on scallop growth in natural habitats. Although the MAB is one of the most well studied shelf ecosystems globally, bottom temperature measurements are still scarce with larger spatiotemporal gaps than those at other depths in the MAB (du Pontavice *et al*., 2023). High-resolution physical models are important complementary bottom temperature data sources and have been used to investigate benthic thermal habitat suitability, groundfish stock assessment, and fish recruitment dynamics in the MAB and surrounding marine ecosystems (Fogarty *et al*., 2008; Tanaka *et al*., 2020; Du Pontavice *et al*., 2022). Although physical models cannot perfectly reproduce realistic bottom thermal conditions due to the challenges in capturing high frequency events, unrealistic bathymetry, grid resolutions, uncertainties in forcing inputs, and many other factors, they can still provide spatially and temporally continuous results with acceptable accuracy (Li *et al*., 2017; du Pontavice *et al*., 2023). To enhance the quality of modeled bottom temperature, observational data are synthesized with model outputs using data assimilation and statistical bias correction methods (Chen *et al*., 2009; Chang *et al*., 2021; du Pontavice *et al*., 2023). These reanalysis products better match in-situ measurements with reduced errors. However, several degree discrepancies between bias-corrected products and observations are still common at those locations with fewer observations, contributing to the uncertainties in thermal stress intensity estimation (Li *et al*., 2017; Chang *et al*., 2021). As an alternative tool for producing gap-free bottom temperature data in sea scallop habitats, machine learning models have been developed rapidly in recent years and applied to reconstruct water temperature at depths (Su *et al*., 2018, 2022; Wang *et al*., 2021). Compared with numerical models, machine learning models have unique advantages in efficiently capturing complex relationships between water temperature and its drivers without compromising the quality of outputs. In future efforts, synthesizing bottom temperature data from multiple sources can help us better constrain the uncertainties in thermal stress estimations and provide valuable information for the development of adaptive fishery management plans under climate warming.

### 4.2 Possible drivers of sea scallop growth interannual variations in the MAB

In this study, we observed broadly synchronized sea scallop growth interannual variations across the four subregions prior to the emergence of growth rate spatial heterogeneity driven by thermal stress (Fig. 4 and Table. 1). Our results imply that scallop growth throughout the MAB was influenced by shared environmental factors before 2015 — likely through a mechanism known as the Moran effect (Moran, 1953). However, the drivers of sea scallop growth rate interannual variation remain poorly understood. We examined several potential variables (e.g., DD in the optimum temperature range between 10 and 15 °C, mean bottom temperature, and fishing mortality in the MAB) but found no significant correlations between these factors and scallop growth rate. Additionally, scallop growth interannual variation in the MAB was not caused by biased sampling because sampling sites and shells for analysis were randomly selected. The interannual variations in averaged sampling depths and mean scallop size were not significantly correlated with scallop growth rates (Fig. S9). The decoupling between growth rate and the variables we examined suggests that scallop growth interannual variation in the MAB before 2015 might be driven by a factor whose magnitude is still difficult to quantify or by the synergistic effects of multiple stressors.

The biotic and abiotic factors modulating sea scallop growth have been extensively investigated in previous studies. In addition to temperature effects we focused on in this work, food condition (i.e., food quantity and quality) is another critical factor limiting sea scallop growth (Pilditch and Grant, 1999a). Laboratory studies show that favorable food supply can facilitate rapid growth of sea scallops (MacDonald and Thompson, 1986; Cranford and Grant, 1990). In the MAB, reduced scallop growth rates in the deep regions are ascribed to limited food supply offshore (Hart and Chute, 2009; Zang *et al*., 2022a). Although food condition has been confirmed as an important factor modulating sea scallop growth rate, quantifying the magnitude of food limitation in natural habitats is still difficult due to several reasons. First, food availability for scallops is jointly influenced by various physical and biogeochemical processes. The primary food items for scallops are phytoplankton and detritus (Shumway *et al*., 1987). Pelagic phytoplankton abundance in the MAB show pronounced variability across multiple spatiotemporal scales (Xu *et al*., 2011, 2020), and its vertical movement due to mixing, advection, and sinking further complicates the relationship between pelagic phytoplankton dynamics and phytoplankton available for scallops in the bottom boundary layer (BBL; Gemmell *et al*., 2016). Additionally, benthic diatoms are also major food sources for scallops, especially for those in the deep water (Shumway *et al*., 1987). Detritus is an important food supplement for sea scallops, and its concentration in the BBL is strongly influenced by hydrodynamics and biogeochemical cycling (e.g., vertical settling, mixing, decomposition, and resuspension) (Munroe *et al*., 2013; Zang *et al*., 2021, 2022a). The complex interactions between physical/biogeochemical processes and scallop food dynamics present significant challenges in determining food condition within the BBL. Second, collecting food information in the BBL remains subject to technical constraints. Most in-situ measurements are limited to near-bottom depths. However, complex hydrodynamics in the BBL contribute to the strong vertical gradient in food concentration near the sediment-water interface, and using near-bottom data as a proxy of food condition probably will introduce large uncertainties in quantifying food condition for scallops in the BBL (Fréchette *et al*., 1989; Newell *et al*., 2005). The lack of field-based scallop food concentration measurements also leads to the difficulties in assessing food condition based on biological model outputs as it hinders the validation of model results in the BBL. Third, scallop feeding activities can be influenced by ambient environments, resulting in the decoupling between food availability and uptake. For example, the optimum current velocity for scallop feeding/growth is around 10 cm/s, and strong or weak tidal currents contribute to lower clearance rate (Wildish and Saulnier, 1992; Cranford *et al*., 1998; Pilditch and Grant, 1999b). High water turbidity due to strong sediment resuspension also inhibit scallop feeding, although detritus entrainment provides extra food for scallops (Grant *et al*., 1997).

In addition to thermal and food conditions, other environmental and anthropogenic factors can also impact scallop growth rates. OA driven by increased oceanic absorption of atmospheric CO_2_ has increased global ocean acidity and is adverse to the growth of calcifying organisms (Saba *et al*., 2019). Although the causal linkage between sea scallop growth rate and OA has not been detected in natural habitats, laboratory experiments and modeling studies have confirmed decreased scallop growth under lower pH levels (Cameron *et al*., 2022; Pousse *et al*., 2023). Parasitism by boring sponges and prokaryotes has long been reported as an important driver of decreased sea scallop growth and mass mortality because more energy is allocated to repairing the damaged structure (Medcof, 1949; Inglis *et al*., 2016; Levesque *et al*., 2016; Siemann *et al*., 2019). Fishing activities can also lower sea scallop growth by selectively removing fast-growing individuals from the population (Hart and Chute, 2009). It is worth noting that the interactive effects of multiple environmental factors may be responsible for the interannual variation in scallop growth. For example, the concurrent warming and OA exacerbate sea scallop growth and mortality by increasing energy consumption to maintain metabolism, while the negative effects of OA can be offset by sufficient food supply to some extent (Sanders *et al*., 2013; Cameron *et al*., 2022; Stechele and Lavaud, 2024). Therefore, in addition to quantifying the magnitudes of various stressors in scallop habitats and their impacts on scallop growth, understanding the phenological patterns of these factors and their synergistic effects on scallop physiology is also critically important in unveiling the drivers of scallop growth interannual variability in the MAB. Such effort requires high-quality environmental stressor datasets with adequate temporal and spatial resolutions.

### 4.3 Implications for scallop resource management under future warming

The implementation of sea scallop management strategies (e.g., limits on days at sea, gear restrictions, and rotational closures of fishing grounds) in the U.S. has successfully restored sea scallop populations and increased the commercial harvest since 1994 (Lee *et al*., 2019). However, warming-induced scallop habitats contraction, growth rate decrease, size structure deterioration, and mass mortality over the last two decades suggest that thermal stress is becoming more severe and threatening the scallop fishery in the MAB. Although our results showed that the subpopulations in the deep MAB benefitted from warming before the thermal condition reached a threshold (Fig. 4), the continued temperature increase will ultimately result in the entire MAB becoming unsuitable for scallop growth and survival. The populations in the northern habitats (e.g., Georges Bank and the Great South Channel) will also be negatively influenced by long-term climate warming (Tanaka *et al*., 2020; Zang *et al*., 2023). In addition, the spatial pattern of thermal stress probably will become more complex in the future: thermal stress in the MAB is overall stronger in the shallow area with low latitude (shallow SMAB in Fig. 5), while the 2022 sea scallop die-off due to warming occurred in the Elephant Trunk in the central part of the MAB scallop habitat (NOAA, 2024). It is still challenging to develop adaptive strategies to mitigate the negative effects of thermal stress on scallop populations because warming events are usually associated with large scale physical processes (e.g., Gulf Stream meandering, warm core ring impingement, and atmospheric jet stream), and scallop thermal acclimation probably cannot keep pace with the warming rate in the MAB (Gawarkiewicz *et al*., 2012; Chen *et al*., 2014). Nevertheless, continued monitoring efforts in the MAB are still essential to provide critical insights into scallop growth and distribution, which are vital for informing commercial fishing activities (e.g., fishing site selection), developing proactive management plans for other scallop habitats, and disentangling the intricate relationships between scallop population dynamics and multi-stressors.

Given the projected decreases in scallop growth rate and biomass under climate warming, sea scallop aquaculture may be an alternative to offset the imbalance between decreased wild harvest and high market demand. Aquaculturists in the Northwest Atlantic began to establish cultured sea scallop industry as early as the 1970s, but its development lags far behind the rest of the world and only contributes to ∼ 0.01% of the total production (Shumway and Parsons, 2016). Several bottlenecks related to slow growth of scallops, biofouling, high natural mortality, site selection, and high costs of labor and equipment hinder the expansion of sea scallop aquaculture in the U.S. (Penney and Mills, 2000; Robinson *et al*., 2016; Shumway and Parsons, 2016). Increased investments in sea scallop aquaculture research and technology over the last decade have overcome some of the aforementioned challenges and provide a valuable knowledge base for the success of sea scallop aquaculture (Grant *et al*., 2003; Coleman *et al*., 2021, 2022; Ishaq *et al*., 2023). Future efforts to develop adaptive, science-based strategies for aquaculture site selection, cultivation, and operations will enhance scallop productivity and complement wild-capture fisheries to mitigate the impacts of warming on sea scallop industry.

## 5. Conclusions

We estimated the spatiotemporal patterns of sea scallop growth rate and thermal stress intensity in the MAB from 2000 to 2018. The results suggested that scallop growth rates across all size groups were lower in the deep subregions, while their latitudinal gradients were much weaker. Scallop growth rates in the entire MAB were generally synchronized until 2014, after which strong thermal stress in 2015 and 2016 coincided with the emerging spatial heterogeneity featured by higher growth rate in the deep subregions and lower growth rate in the shallow areas, respectively. A dynamic energy budget model was adopted to assess sea scallop growth under 2016 and climatological thermal conditions. The comparisons between model results and observational data indicated the good performance of DEB model in capturing scallop growth dynamics in the MAB. The DEB model results suggested that bottom temperature exceeding 16 °C in 2016 profoundly lowered energy allocation to scallop growth in the shallow subregions, while warming in the deep subregions lead to more favorable thermal conditions for scallop growth. Given the projected future warming in most sea scallop habitats on the Northeast U.S. Shelf, using appropriate metrics to quantify thermal stress intensity and addressing the joint effects of multiple stressors on scallop growth are critical for understanding the drivers of sea scallop growth rate variations and developing adaptive management strategies to sustain sea scallop fishery resources.

## Acknowledgements

This study was supported by NSF Northeast US Shelf Long-Term Ecological Research (NES-LTER) Program (OCE-2322676), and NOAA Sea Scallop Research Set-Aside (RSA) Program (NA22NMF4540060 and NA24NMFX454G0012) in collaboration with Cape Cod Commercial Fishermen’s Alliance, and also RSA program (NA22NMF4540046, and NA22NMFX454G007 T1-01).

## Author contributions

Z.Z. and R.J. designed the study and performed the statistical analyses. Z.Z. and R.L. developed and calibrated the DEB model. C.C. and S.L. provided physical model setup information and outputs. R.J. and D.R.H. assisted with the conceptualization of this work. Z.Z., R.J., D.R.H., and R.L. developed the first manuscript draft. All authors reviewed the paper and approved the manuscript submission.

## Data availability

The sea scallop growth rate data and the DEB model outputs used in this article will be shared upon a reasonable request to the corresponding author.

