## Supporting Text, Figs S1-S9, and Tables S1-S2 for "Spatially variable growth responses to warming in Atlantic Sea Scallops (*Placopecten magellanicus*)": Supplementary_Material_submission.pdf

### Supporting Text: growth rate anomaly analysis

We estimated growth rate anomaly for each sample using field measurement and von Bertalanffy equation:

$$L_{yr+1} = \exp(-K)\, L_{yr} + L_{\infty}[1 - \exp\,(-K)] \tag{1}$$

Where $K = 0.574 - 0.0012$ depth and $L_{\infty} = 309.0 - 0.947\text{depth} - 23.1\text{lat}$ (Table 2 in Hart and Chute, 2009). For each observed growth increment $L_{yr}$, modeled scallop shell height after one year ($L_{yr+1,model}$) is estimated using equation 1 and its corresponding depth and latitude. Modeled scallop shell increment $I_{yr+1,model}$ equals to $L_{yr+1,model} - L_{yr}$. The scallop growth rate anomaly $A$ is estimated using equation (2):

$$A = \frac{I_{yr+1,obs} - I_{yr+1,model}}{I_{yr+1,model}} \tag{2}$$

Since $I_{yr+1,model}$ already have depth and latitude effects built-in, the anomaly $A$ is independent of depth and latitude. In each subregion, annual mean growth rate is estimated by averaging all the anomalies in the corresponding year (Fig. S5).

**Sample collected in year Y**

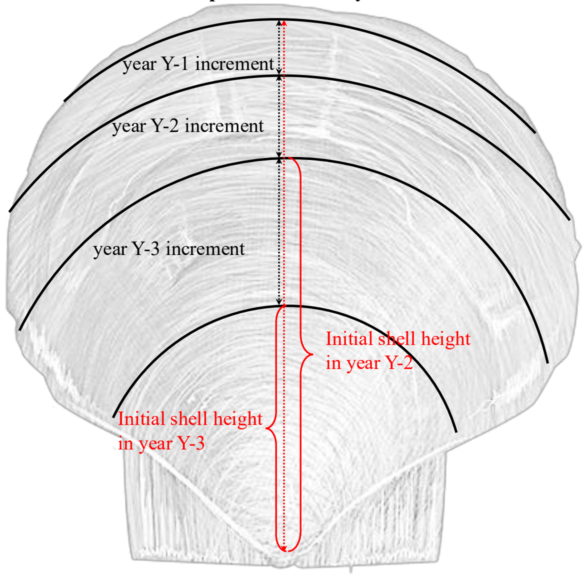


Fig. S1. Scallop growth diagram showing the increments between rings (black lines) and their corresponding years. The two red braces are the examples showing initial shell height at the beginning of year Y-3 and year Y-2.

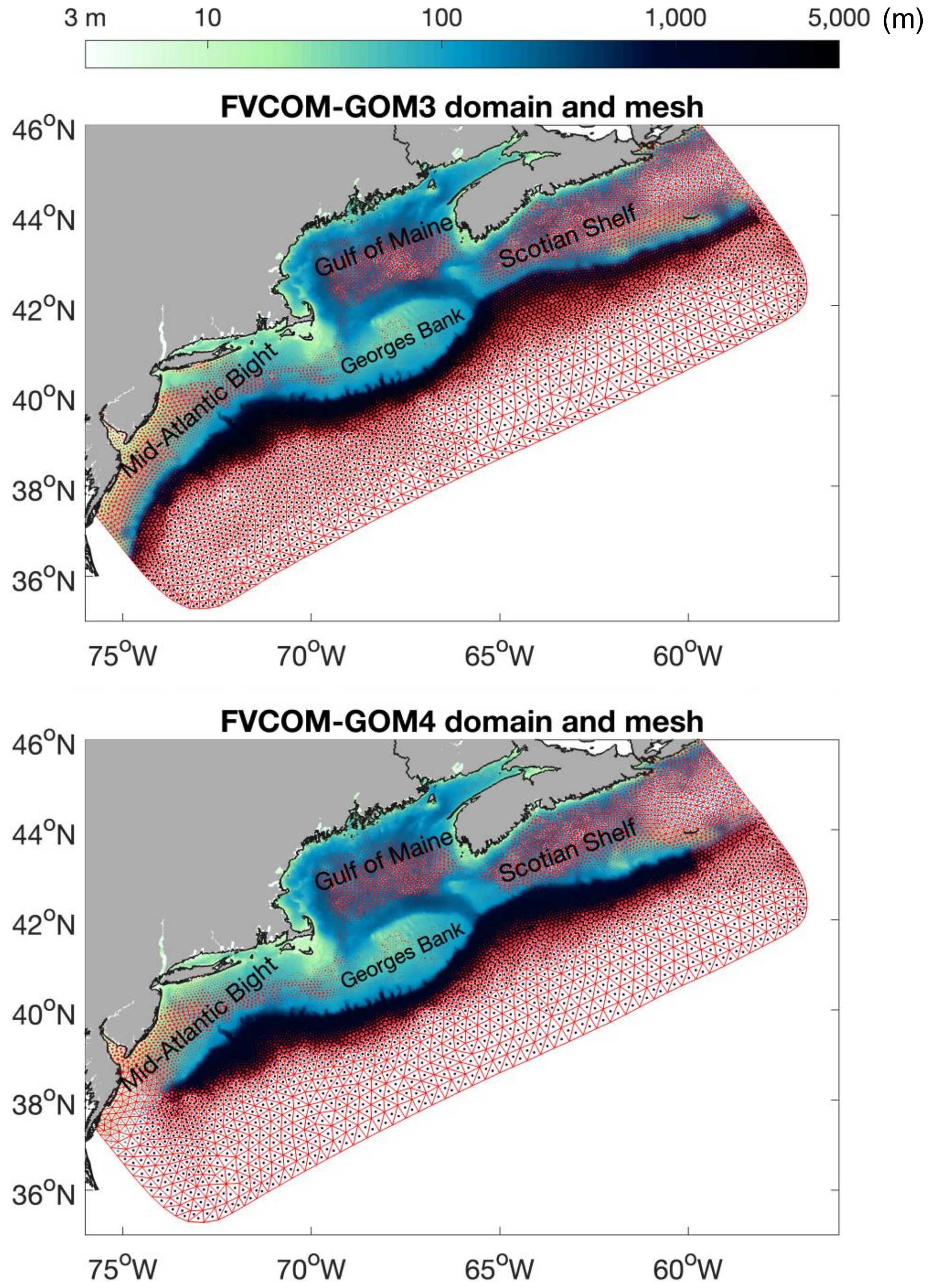


Fig. S2. Model domains of the FVCOM-GOM3 (upper panel) and the FVCOM-GOM4 (lower panel) overlaid with water depth.

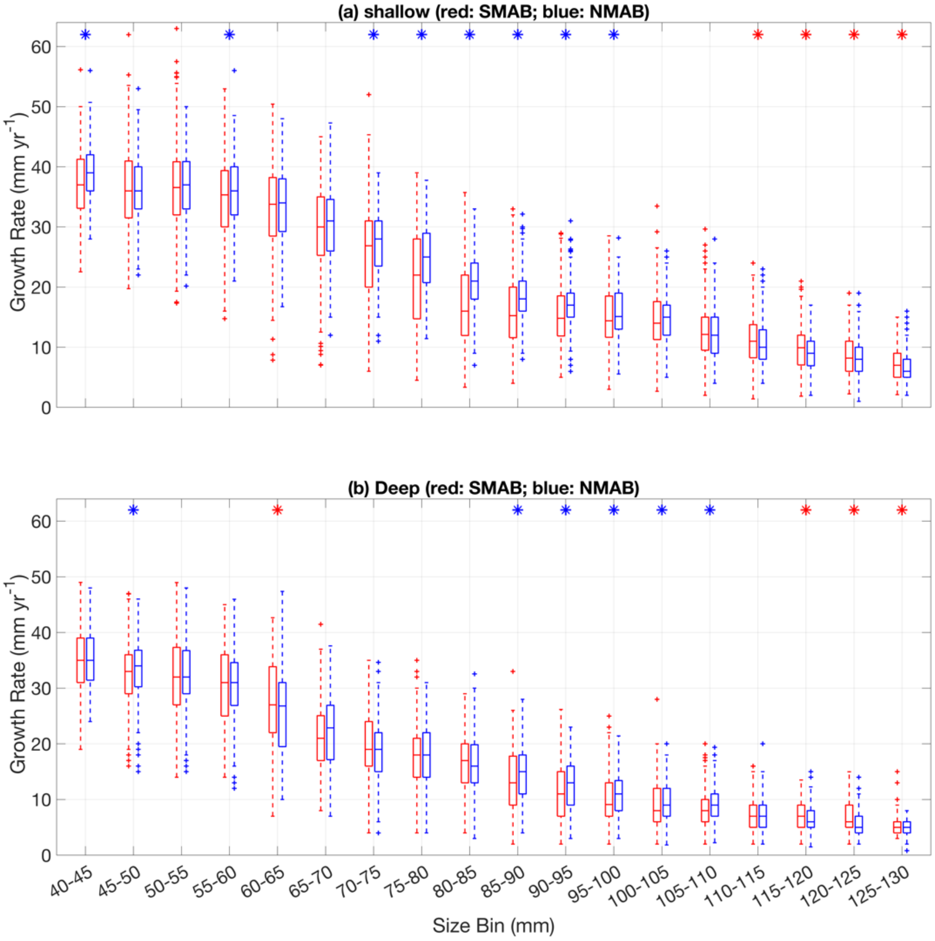


Fig. S3. Scallop growth rates across 18 size bins (x-axis) in the shallow (a) and deep (b) subregions. Red and blue colors indicate data in the SMAB and the NMAB subregions, respectively. Mann–Whitney U tests were performed for each size bin to assess statistical significance. Red and blue asterisks indicate higher values in the SMAB and the NMAB subregions, respectively.

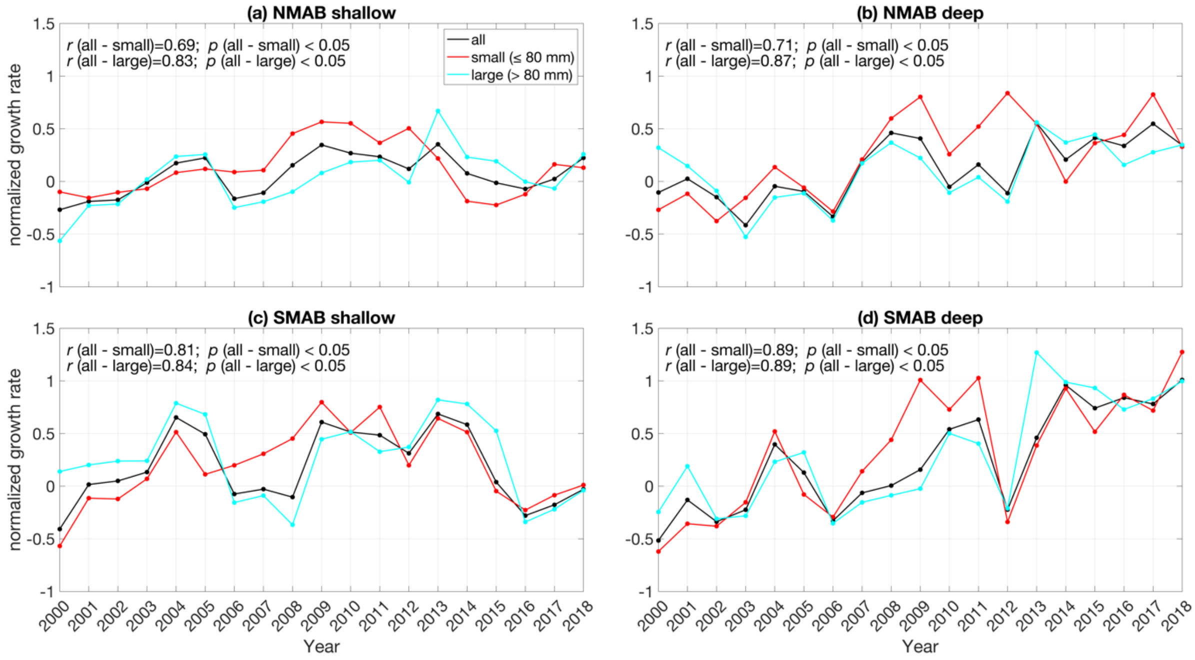


Fig. S4. Interannual variations in normalized sea scallop growth rates for small size groups (≤ 80 mm; red), large size groups (> 80 mm; cyan), and the entire size groups (black; same as the black lines in Fig. 4) in the four subregions.

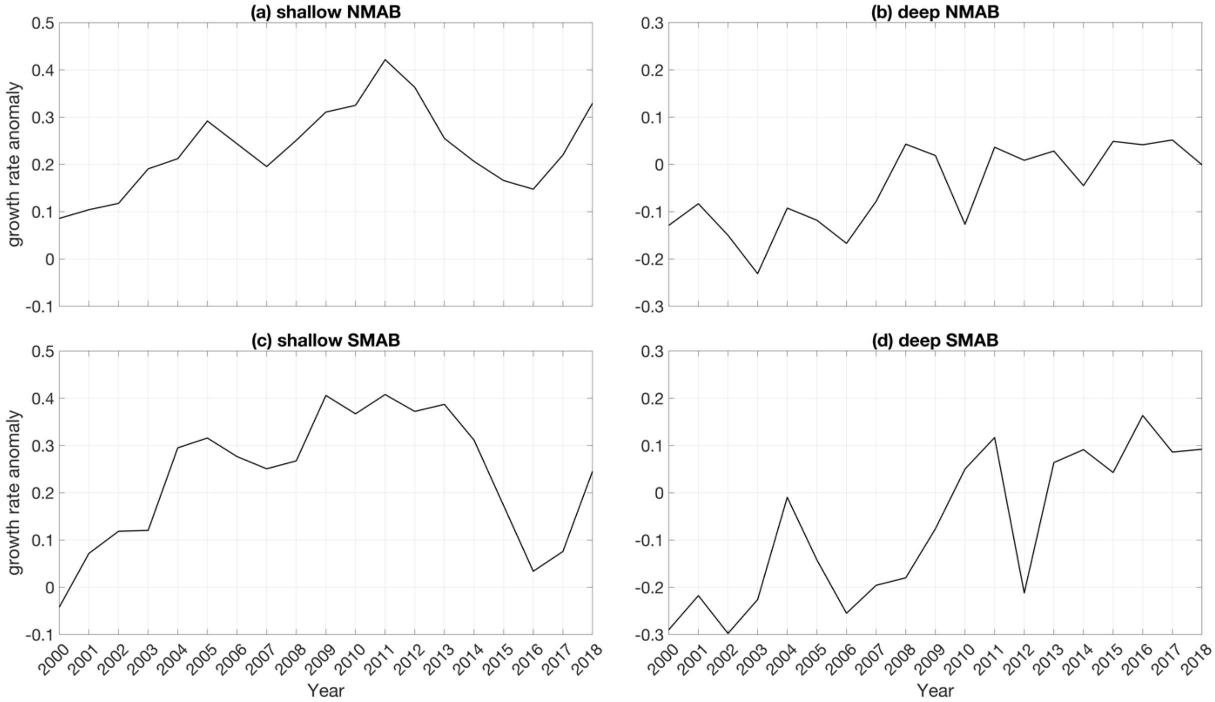


Fig. S5. Interannual variations in mean sea scallop growth rate anomalies in the four subregions.

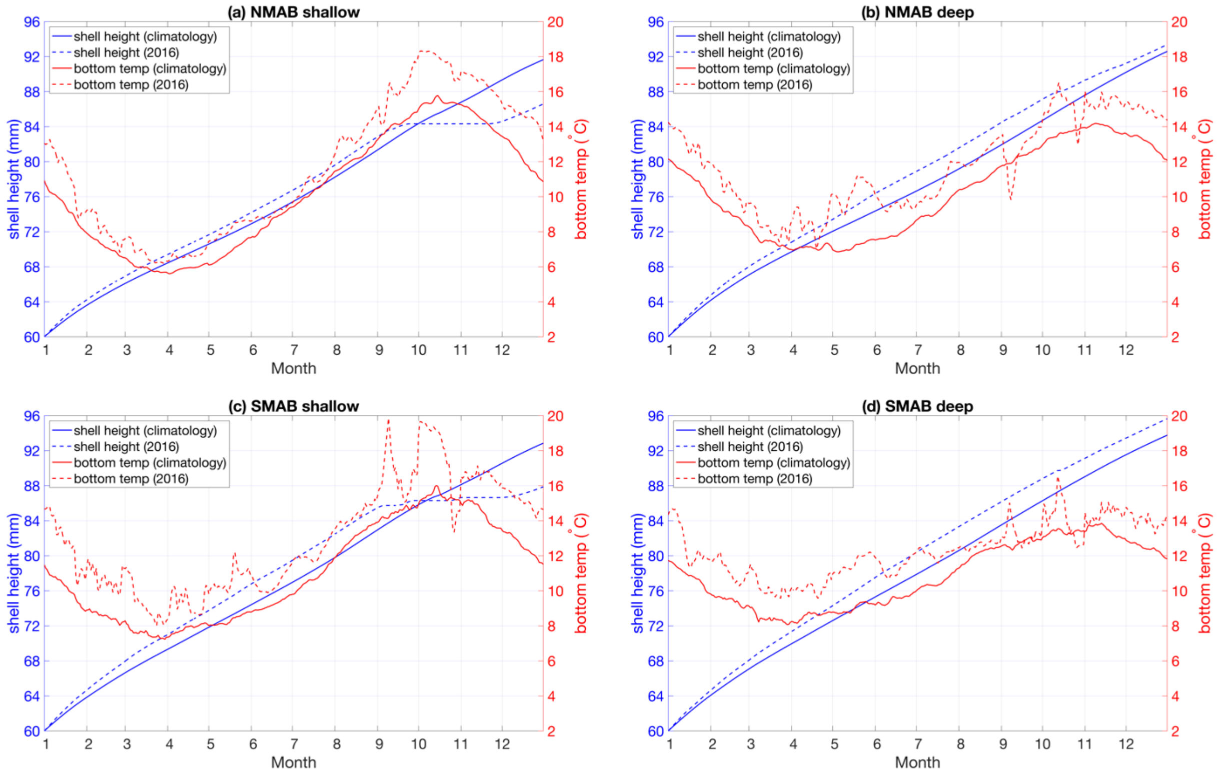


Fig. S6. Annual shell height growth (blue) and the corresponding bottom temperature (red) in the four subregions (panel a: shallow NMAB; panel b: deep NMAB; panel c: shallow SMAB; panel d: deep SMAB). The initial scallop shell height is assigned as 60 mm. Solid and dashed lines are the results based on bottom temperature climatology and bottom temperature in 2016, respectively.

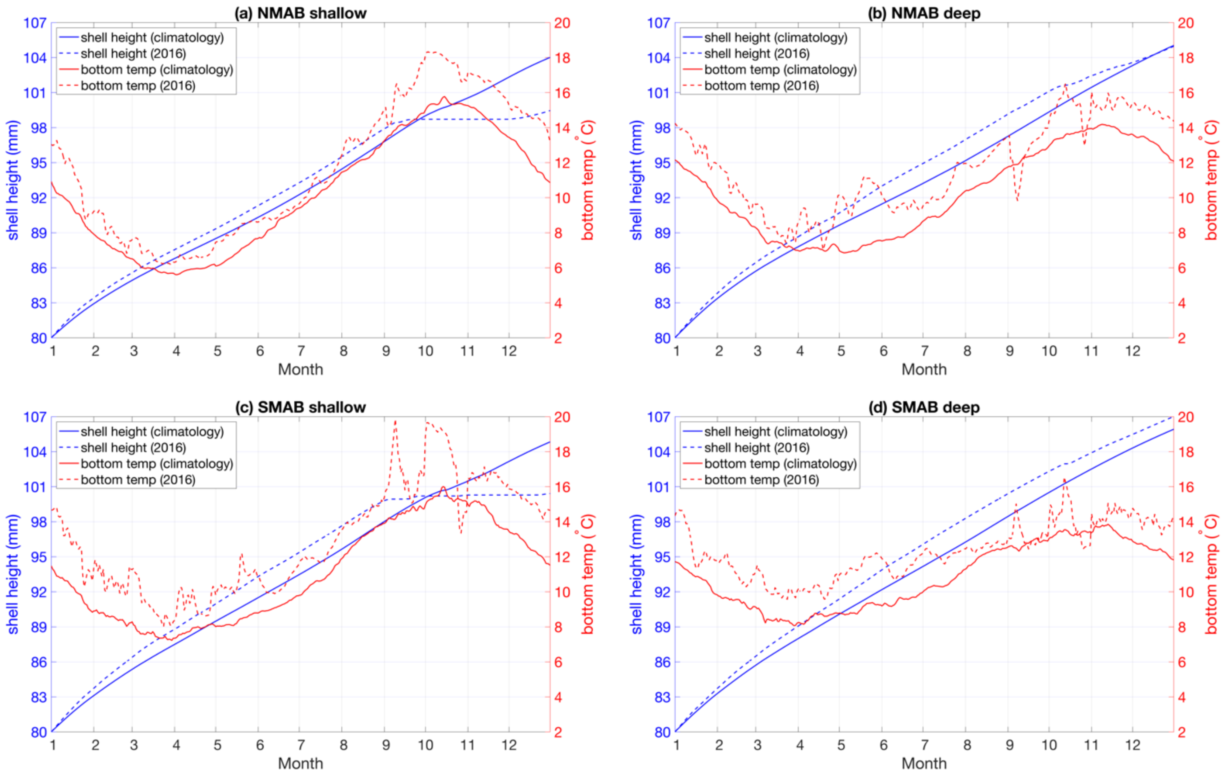


Fig. S7. Same as Fig. S5 but with an initial shell height of 80 mm.

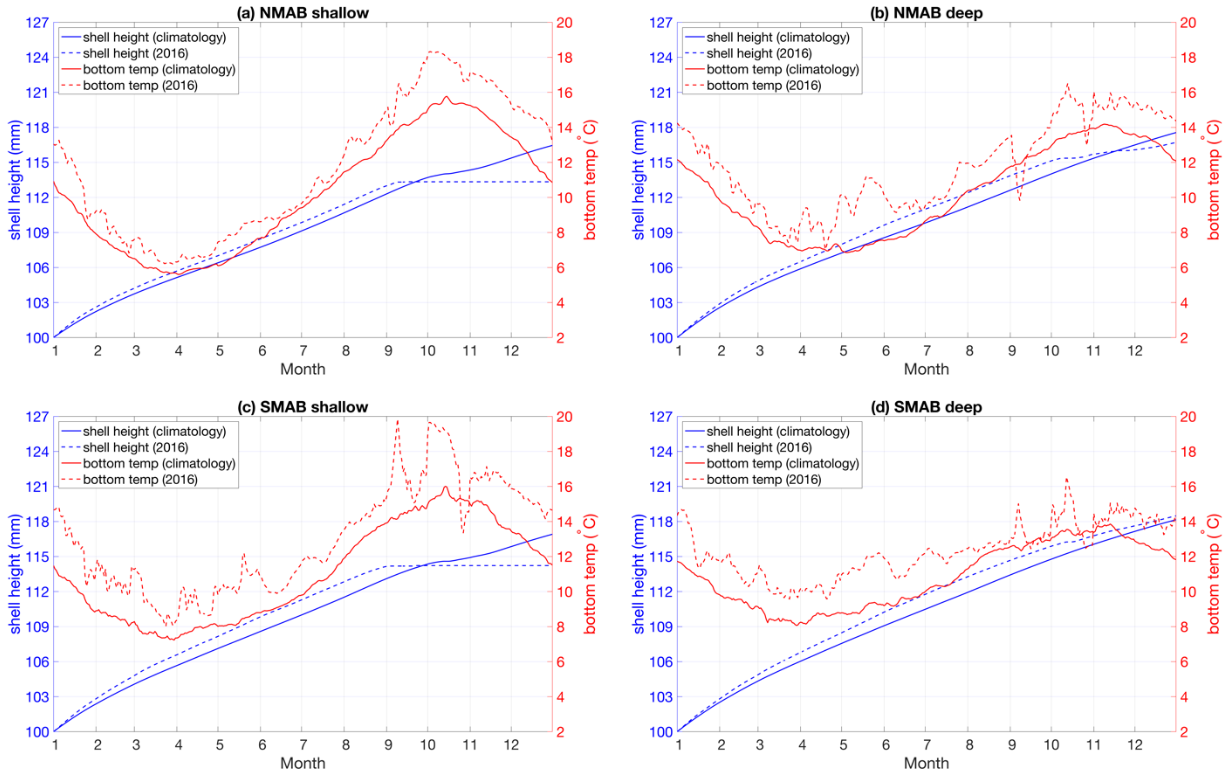


Fig. S8. Same as Fig. S5 but with an initial shell height of 100 mm.

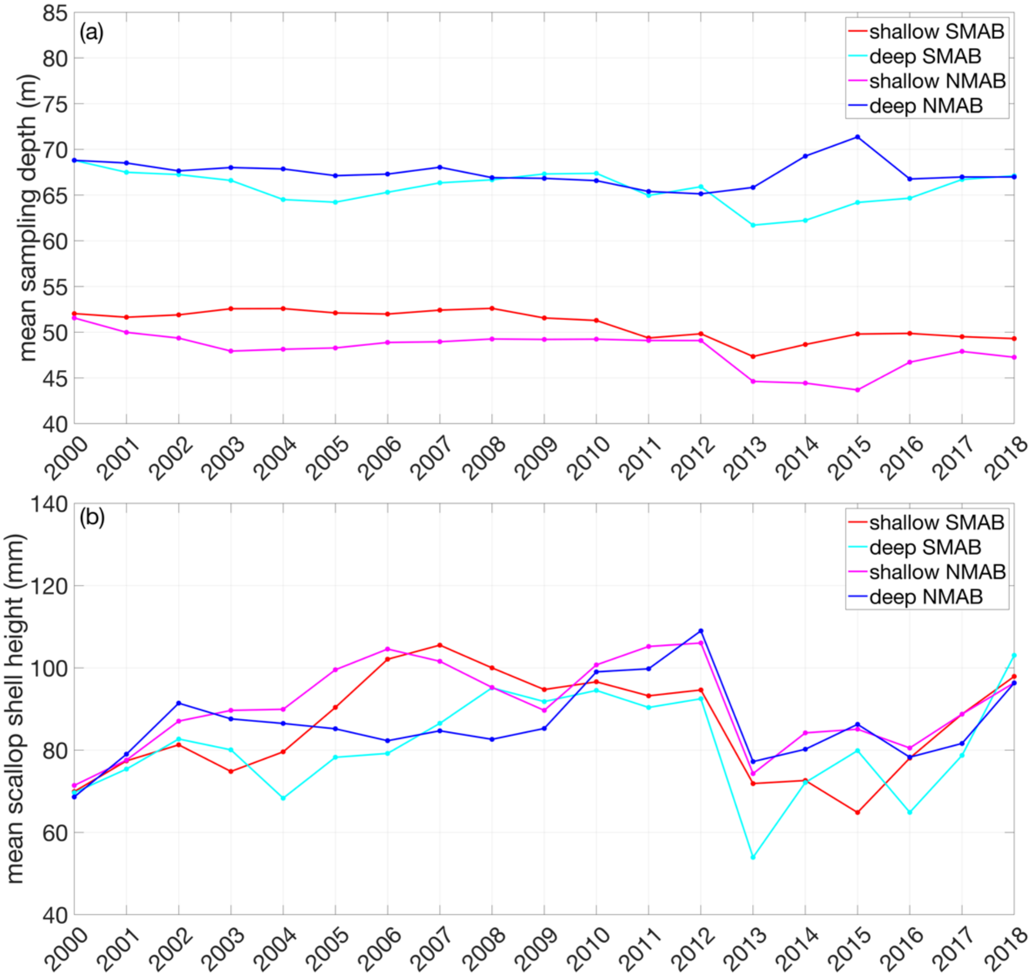


Fig. S9. Interannual variations in mean sampling depth (a) and sampled scallop shell height (b) in the four subregions.

Table S1. DEB model variables and equations.

| Energy Fluxes Equations | Model Dynamics/Outputs |
|---|---|
| Assimilation Flux: $\dot{p}_A = \dot{P}_{Am} f V^{2/3} c_{T,G}$ | Temperature Correction Function for Growth: $$c_{T,G} = \exp\left(\frac{T_A}{T_{ref,G}} - \frac{T_A}{T}\right) \cdot \frac{[1+\exp\left(\frac{T_{AL}}{T} - \frac{T_{AL}}{T_{L,G}}\right) + \exp\left(\frac{T_{AH}}{T_{H,G}} - \frac{T_{AH}}{T}\right)]^{-1}}{[1+\exp\left(\frac{T_{AL}}{T_{ref,G}} - \frac{T_{AL}}{T_{L,G}}\right) + \exp\left(\frac{T_{AH}}{T_{H,G}} - \frac{T_{AH}}{T_{ref,G}}\right)]^{-1}}$$ |
| Mobilization Flux: $\dot{p}_C = E\frac{[E_G]V^{2/3}\dot{v}c_{T,G} + \dot{p}_M}{\kappa E + [E_G]V}$ | Temperature Correction Function for maintenance: $$c_{T,R} = \exp\left(\frac{T_A}{T_{ref,R}} - \frac{T_A}{T}\right) \cdot \frac{[1+\exp\left(\frac{T_{AL}}{T} - \frac{T_{AL}}{T_{L,R}}\right) + \exp\left(\frac{T_{AH}}{T_{H,R}} - \frac{T_{AH}}{T}\right)]^{-1}}{[1+\exp\left(\frac{T_{AL}}{T_{ref,R}} - \frac{T_{AL}}{T_{L,R}}\right) + \exp\left(\frac{T_{AH}}{T_{H,R}} - \frac{T_{AH}}{T_{ref,R}}\right)]^{-1}}$$ |
| Somatic Maintenance Flux: $\dot{p}_M = [\dot{P}_M] c_{T,R} V$ | Structural Volume: $\frac{dV}{dt} = \frac{\dot{p}_G}{[E_G]} - \dot{p}_{LV}$ |
| Allocation Flux to Growth: $\dot{p}_G = \max(0, \kappa\dot{p}_C - \dot{p}_M)$ | Reserve: $\frac{dE}{dt} = \dot{p}_A - \dot{p}_C$ |
| Maturity Maintenance Flux: $\dot{p}_J = E_H^p k_J c_{T,R}$ | Reproduction Buffer: $\frac{dE_R}{dt} = \dot{p}_R\kappa_R - \dot{p}_{Go} - \dot{p}_{LR}$ |
| Allocation Flux to Reproduction: $\dot{p}_R = \max(0, (1-\kappa)\dot{p}_C - \dot{p}_J)$ | Energy in Gonads: $\frac{dE_{Go}}{dt} = \dot{p}_{Go} - E_{SP}$ |
| Allocation Flux to Gonads: $\dot{p}_{Go} = \kappa_R E_R (\frac{\dot{v}c_{T,G}}{V^{\frac{1}{3}}} + \frac{[\dot{P}_M]c_{T,R}}{[E_G]})(1 - \kappa\frac{E}{[E_G]V + \kappa E})$, if $gsi > gsi_{sp}$ | Shell Height: $SH = \frac{V^{\frac{1}{3}}}{\delta_M}$ |
| Spawned Energy: $E_{SP} = \frac{E_{Go}\kappa_R}{dt}$ | Dry Gonad Weight: $DW_{Go} = E_{Go} w_E / \mu_E$ |
| Maintenance Deficit: $Md = \max(0, \dot{p}_M + \dot{p}_J - \dot{p}_C)$ | Wet Gonad Weight: $WW_{Go} = DW_{Go} / d_{V\_gonad}$ |
| Lysis of Reproduction Buffer: $\dot{p}_{LR} = \min(Md, \frac{E_R\kappa_R}{dt})$ | Dry Tissue Weight: $DW_{Tissue} = V d_{V_{somatic}} + \frac{(E+E_R)w_E}{\mu_E} + DW_{Go}$ |
| Lysis of Structure: $\dot{p}_{LV} = \min(Md - \dot{p}_{LR}, \frac{V d_V \mu_V}{w_V dt})$ | Wet Tissue Weight: $WW_{Tissue} = DW_{Tissue} / d_{V_{somatic}}$ |

Table. S2. Sea scallop DEB model parameters

| Parameter Description | Symbol | Value | Unit | Source |
|---|---|---|---|---|
| maximum surface area-specific assimilation rate | $\dot{P}_{Am}$ | 396 | J d$^{-1}$ cm$^{-2}$ | 1 |
| energy conductance | $\dot{v}$ | 0.0065 | cm d$^{-1}$ | This study |
| fraction of energy allocated to soma | $\kappa$ | 0.76 | — | 1 |
| reproduction efficiency | $\kappa_R$ | 0.95 | — | 1 |
| volume-specific somatic maintenance rate | $[\dot{P}_M]$ | 311 | J d$^{-1}$ cm$^{-3}$ | 1 |
| maturity maintenance rate | $k_J$ | 0.002 | d$^{-1}$ | 1 |
| volume-specific cost for structure | $[E_G]$ | 2393 | J cm$^{-3}$ | 1 |
| maturity at puberty | $E_H^p$ | 863 | J | 1 |
| shape coefficient | $\delta_M$ | 0.15 | — | 1 |
| chemical potential of structure | $\mu_V$ | 500,000 | J mol$^{-1}$ | 1 |
| chemical potential of reserve | $\mu_E$ | 550,000 | J mol$^{-1}$ | 1 |
| molecular weight of structure | $w_V$ | 23.9 | g mol$^{-1}$ | 1 |
| molecular weight of reserve | $w_E$ | 23.9 | g mol$^{-1}$ | 1 |
| gonado-somatic index threshold for spawning | $gsi_{sp}$ | 0.04 | — | 2,3 |
| food limiting factor | $f$ | 0.64 | — | This study |
| Arrhenius temperature | $T_A$ | 5290 | K | 4 |
| Arrhenius temperature for lower boundary | $T_{AL}$ | 4621 | K | 1 |
| Arrhenius temperature for upper boundary | $T_{AH}$ | 94252 | K | 1 |
| Reference temperature for growth | $T_{ref,G}$ | 279 | K | 5 |
| Reference temperature for respiration | $T_{ref,R}$ | 293 | K | 1 |
| Lower boundary tolerance range for growth | $T_{L,G}$ | 275 | K | 1 |
| Lower boundary tolerance range for respiration | $T_{L,R}$ | 275 | K | 4 |
| Upper boundary tolerance range for growth | $T_{H,G}$ | 290 | K | 1 |
| Upper boundary tolerance range for respiration | $T_{H,R}$ | 297 | K | 6 |


[1] Lavaud, R., and Kooijman, S. A. L. M. 2020. AmP Placopecten magellanicus, version 2020/03/21.

[2] Thompson, K. J., Inglis, S. D., and Stokesbury, K. D. E. 2014. Identifying spawning events of the sea scallop placopecten magellanicus on Georges Bank. Journal of Shellfish Research, 33: 77–87.

[3] Sarro, C. L., and Stokesbury, K. D. E. 2009. Spatial and Temporal Variation in the Shell Height/Meat Weight Relationship of the Sea Scallop Placopecten magellanicus in the Georges Bank Fishery. Journal of Shellfish Research, 28: 497–503.

[4] Van der Veer, H. W., Cardoso, J. F. M. F., and Van der Meer, J. 2006. The estimation of DEB parameters for various Northeast Atlantic bivalve species. Journal of Sea Research, 56: 107–124.

[5] Grant, J., and Cranford, P. J. 1991. Carbon and nitrogen scope for growth as a function of diet in the sea scallop Placopecten magellanicus. Journal of the Marine Biological Association of the United Kingdom, 71: 437–450.

[6] Zang, Z., Ji, R., Hart, D. R., Chen, C., Zhao, L., and Davis, C. S. 2022. Modeling Atlantic sea scallop (Placopecten magellanicus) scope for growth on the Northeast U.S. Shelf. Fisheries

Oceanography, 31: 271–290.
